# Origin and recent expansion of an endogenous gammaretroviral lineage in canids

**DOI:** 10.1101/414607

**Authors:** Julia V. Halo, Amanda L. Pendleton, Abigail S. Jarosz, Robert J. Gifford, Malika L. Day, Jeffrey M. Kidd

## Abstract

Mammalian genomes contain a fossilized record of ancient retroviral infections in the form of endogenous retroviruses (ERVs). We used whole genome sequence data to assess the origin and evolution of the recently active ERV-Fc gammaretroviral lineage based on the record of past infections retained in the genome of the domestic dog, *Canis lupus familiaris.* We identified 165 loci, including 58 insertions absent from the dog reference assembly, and characterized element polymorphism across 332 canids from nine species. Insertions were found throughout the dog genome including within and near gene models. Analysis of 19 proviral sequences identified shared disruptive mutations indicating defective proviruses were spread via complementation. The patterns of ERV polymorphism and sequence variation indicate multiple circulating viruses infected canid ancestors within the last 20 million to within 1.6 million years with a recent bust of germline invasion in the lineage leading to wolves and dogs.

## Introduction

During a retroviral infection, the viral genome is reverse transcribed and the resulting DNA is then integrated into the host genome as a provirus. In principle, the provirus carries all requirements necessary for its replication, and typically consists of an internal region encoding the viral genes (*gag pro*/*pol*, and *env*) flanked by two regulatory long terminal repeats (LTRs) that are identical at the time of integration. Outermost flanking the provirus are short, 4-6 bp target site duplications (TSDs) of host genomic sequence generated during integration. Infection of such a virus within a germ cell or germ tissue may lead to an integrant that is transmitted vertically to offspring as an endogenous retrovirus (ERV). Over time, the ERV may reach detectable frequencies within a population or even fixation within a species (Boeke and Stoye, 1997). Through repeated germline invasion and expansion over millions of years, ERVs have accumulated to such an extent that they account for considerable proportions of genetic sequence of many mammalian genomes.

ERVs have been commonly referred to as ‘fossils’ of their once-infectious counterparts, providing a record of exogenous retroviruses that previously infected a species (Boeke and Stoye, 1997). Because an ERV switches from a relatively rapid evolutionary state as an infectious virus to a relatively slow one while replicated as part of the host genome, recently formed ERVs tend to bear close resemblance to their exogenous equivalent and possess a greater potential to retain functional properties. Across species, the majority of ERVs are thought to provide no advantage to the host, and have, for the most part, been progressively degenerated over time due to accumulated mutations or from recombination between the proviral LTRs that replaces the full-length sequence with a solitary LTR, or ‘solo LTR’ (Boeke and Stoye, 1997). However, increasing evidence suggests evolutionary roles in host physiology via gene regulation, for example by providing alternative promoters, enhancers, splice sites, or termination signals (Chuong et al., 2016, Macfarlan et al., 2012, Rebollo et al., 2012). There are also instances in which ERV gene products have been co-opted for various host functions. Notable examples include syncytial trophoblast fusion in eutherian animals (Lavialle et al., 2013) and blocking of infection from exogenous viruses (Nethe et al., 2005, Stoye, 2012, Blanco-Melo et al., 2017, Weiss and Stoye, 2013).

In humans, ERVs (HERVs) make up over 8% of the genome, the majority being the degenerate remnants of ancient infections. However, a subset of these elements is relatively intact and displays signatures of relatively recent germline invasion. Specifically, the HERV-K (HML-2) group has many young, insertionally polymorphic integrants that display variability in prevalence among global populations, retain ORFs, and includes many copies with high LTR-LTR identity (Wildschutte et al., 2016). Other species are known to harbor similar ‘young’ integrants resulting from relatively recent endogenization events that segregate as unfixed alleles within the species. Examples include the cervid gammaretrovirus in mule deer populations of North America (Elleder et al., 2012) and the insertionally polymorphic ERVs found in domestic and wild cats (Roca et al., 2004, Troyer et al., 2004). Other species are hosts to infection from exogenous viruses that have been shown to contribute to new germline infections, for example, the Koala retrovirus (KoRV) is in the midst of transitioning to an endogenous state in Australia (Ishida et al., 2015, Tarlinton et al., 2006, Lober et al., 2018). Recombination between distinct ERV RNAs that are co-packaged in the same virion may also contribute to new viruses that have altered pathogenic properties (Stocking and Kozak, 2008).

The endogenous retroviruses classified as ERV-Fc are distant relatives of extant gammaretroviruses (also referred to as gamma-like, or γ-like). As is typical of most ERV groups, ERV-Fc is named for its use of a primer binding site complementary to the tRNA used during reverse transcription (tRNA^phe^). Previous analysis of the *pol* gene showed that ERV-Fc elements form a monophyletic clade with the human γ-like ERV groups HERV-H and HERV-W (Jern 2005). As is common to all γ-like representatives, members of the ERV-Fc group possess a ‘simple’ genome that encodes the canonical viral genes and lacks apparent accessory genes that are present among complex retroviruses. The ERV-Fc group was first characterized as a putatively extinct, low copy number lineage that first infected the ancestor of all simians and later contributed to independent germline invasions in primate lineages (Benit et al., 2003). It has since been shown that ERV-Fc related lineages were infecting mammalian ancestors as early as 30 million years ago and subsequently circulated and spread to a diverse range of hosts, including carnivores, rodents, and primates (Diehl et al., 2016). The spread of the ERV-Fc lineage included numerous instances of cross-species jumps and recombination events between different viral lineages, now preserved in the fossil record of their respective host genomes (Diehl et al., 2016).

In comparison to humans and other mammals, the dog (*Canis lupus familiaris*) displays a substantially lower ERV presence, with only 0.15% of the genome recognizably of retroviral origin (Lindblad-Toh et al., 2005, Martinez Barrio et al., 2011). No exogenous retrovirus has been confirmed in the dog or any other canid, though there have been reports of retrovirus-like particles and enzyme activities in affected tissues of lymphomic and leukemic dogs (Ghernati et al., 2000, Modiano et al., 2005, Modiano et al., 1995, Onions, 1980, Perk et al., 1992, Safran et al., 1992, Tomley et al., 1983). Nonetheless, the ERV fossil record in the canine genome demonstrates that retroviruses did infect canine ancestors. The vast majority of canine ERVs (or ‘CfERVs’) are of ancient origin, as inferred by sequence divergence and phylogenetic placement (Martinez Barrio et al., 2011), suggesting most CfERV lineages ceased replicating long ago. An exception comes from a minor subset of ERV-Fc-derived proviruses within the reference genome that possess signatures of recent integration, including high LTR nucleotide identity and the presence of ORFs (Martinez Barrio et al., 2011). This ERV lineage has been recently detailed by Diehl, *et al.*, in which the authors described a distinct ERV-Fc lineage in the Caniformia suborder (Figure 1) classified therein as ‘ERV-Fc1’ (Diehl et al., 2016). The ERV-Fc1 lineage first spread to members of the Caniformia at least 20 million years ago (mya) as a recombinant virus of two otherwise distantly related γ-like lineages: the virus possessed ERV-Fc *gag pol*, and LTR segments but had acquired an *env* gene most closely related to ERV-W (syncytin-like) (Diehl et al., 2016). A derived sublineage, CfERV-Fc1(a), later spread to and infected canid ancestors via a cross-species transmission from an unidentified source, after which the lineage remained active/mobile and endogenized canid members until at least the last 1-2 million years (Diehl et al., 2016). Phylogenetic analyses confirmed the few recently inserted loci belong to CfERV-Fc1(a) (Diehl et al., 2016).

**Figure 1.**
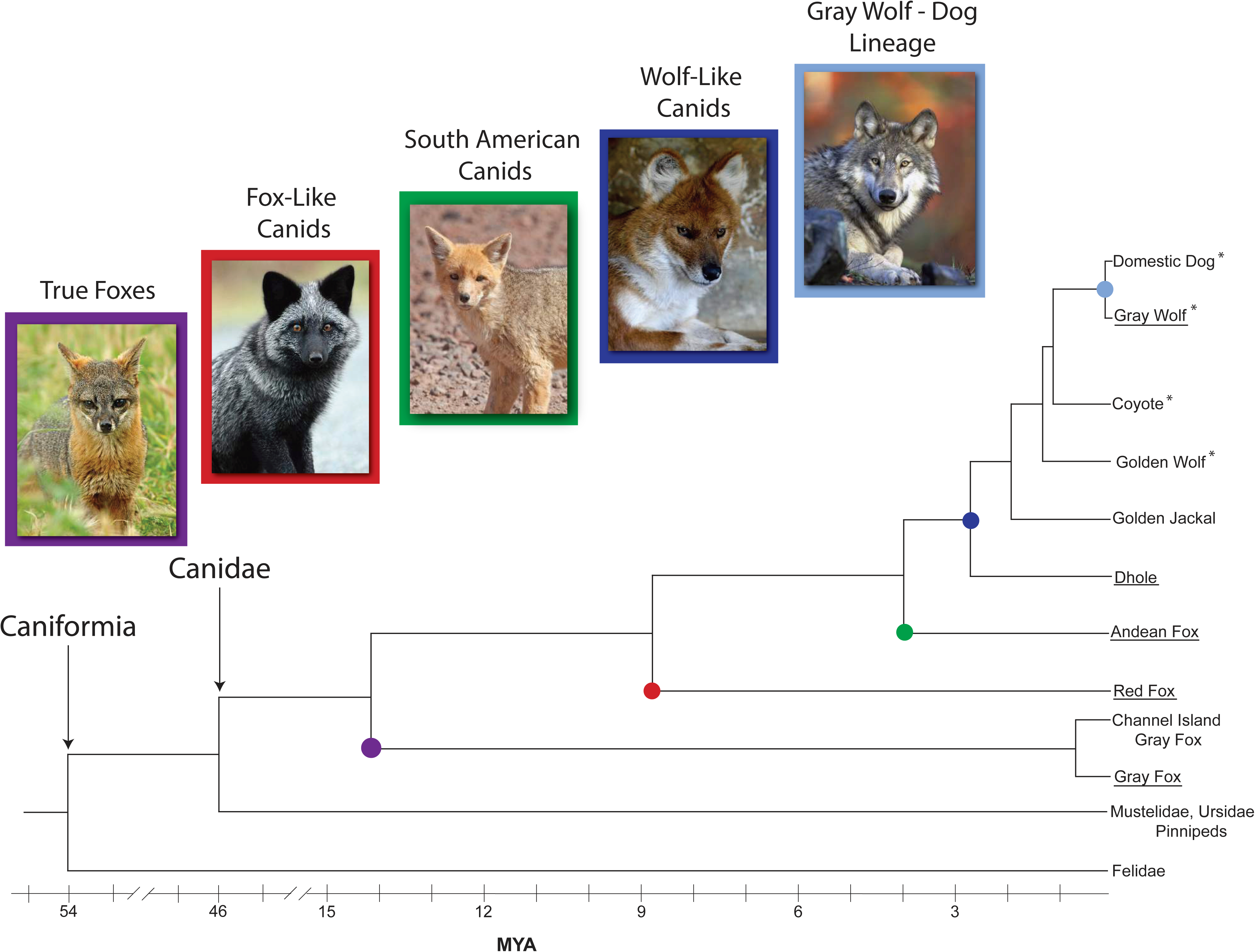
Canidae evolution and representative extant species. Relative to other Caniforms, the evolutionary relationship of the four major canid lineages, along with estimated split times (determined from (Kumar et al., 2017) and (Koepfli et al., 2015)) is shown. Species with asterisks were included in CfERV-Fc1(a) discovery, and all canids here were used for *in silico* genotyping. Images are provided for the underlined species. See acknowledgements for all image credits.

The domestic dog belongs to the family *Canidae*, the oldest family of Carnivora, which arose in North America during the late Eocene (∼46 mya) (Macdonald and Sillero-Zubiri, 2004, Kumar et al., 2017) (Figure 1). Following multiple crossings of the Bering Strait land bridge to Eurasia, canids underwent massive radiations, leading to the ancestors of most modern canids (Macdonald and Sillero-Zubiri, 2004). The now extinct progenitors of the wolf-like canids, belonging to the genus *Canis*, first appeared in North America ∼6 mya and also entered Eurasia via the same route (Macdonald and Sillero-Zubiri, 2004). Slowly, canids colonized all continents excluding Antarctica, as the formation of the Isthmus of Panama permitted dispersal and radiations within South America starting around 3 mya (Macdonald and Sillero-Zubiri, 2004). Approximately 1.1 mya, *Canis lupus*, the direct ancestor of the dog, emerged in Eurasia (Koepfli et al., 2015). Along with many other canid species, the gray wolf migrated back to the New World during the Pleistocene when the land bridge formed once more (Macdonald and Sillero-Zubiri, 2004). Placed within the context of CfERV-Fc1(a) evolution, the initial insertions from this lineage would have occurred while early *Canidae* members were still in North America, and continued until the emergence of the gray wolf.

Utilizing genome data from canid species representing all four modern lineages of Canidae (Figure 1), we assessed the origin, evolution, and impact of the recently active γ-like CfERV-Fc1(a) lineage, yielding the most comprehensive assessment of ERV activity in carnivores to date. Aside from analyses utilizing the current dog reference assembly (CanFam3.1), relatively little is known in this regard. We used Illumina sequence data to characterize CfERV-Fc1(a) integrants in dogs and wild canids, resulting in the discoveries of numerous insertionally polymorphic and novel copies and the further delineation of the presence of this ERV group by comparison of orthologous insertions across species to provide a rich evolutionary history of CfERV-Fc1(a) activity among the *Canidae*. Our analysis demonstrates that the spread of CfERV-Fc1(a) contributed to numerous germline invasions in the ancestors of modern canids, including proviruses with apparently intact ORFs and other signatures of recent integration. The data suggest mobilization of existing ERVs by complementation had a significant role in the proliferation of the CfERV-Fc1(a) lineage in canine ancestors.

## Results

### Discovery of CfERV-Fc1(a) insertions

#### Insertionally polymorphic CfERV-Fc1(a) loci in dogs and wild canids

We determined the presence of CfERV-Fc1(a) insertions using Illumina whole genome sequencing data from dogs and other *Canis* representatives in two ways (Figure 2). First, we searched for CfERV-Fc1(a) sequences in the dog reference genome that were polymorphic across a collection of resequenced canines. In total, our dataset contained 136 CfERV-Fc1(a) insertions, and was filtered to a curated set of 107 intact or near-intact loci, including two loci related by segmental duplication (see Methods). These insertions are referred to as ‘reference’ throughout the text due to their presence in the dog reference genome. Comparative BLAT searches demonstrated their absence from the draft genomes of other extant Caniformia species (*i.e*., ferret and panda). We then intersected the reference loci with deletions predicted by Delly (Rausch et al., 2012) within a sample set of 101 resequenced *Canis* individuals, specifically including jackals, coyotes, gray wolves, and dogs (Table S1). Candidate deletions were classified as those that intersected with annotated ‘CfERVF1’-related loci and were within the size range of the solo LTR or provirus (∼457 and ∼7,885 bp, respectively; Figure 2A). The analysis identified 11 unfixed reference insertions, including 10 solo LTRs and one full-length provirus.

**Figure 2.**
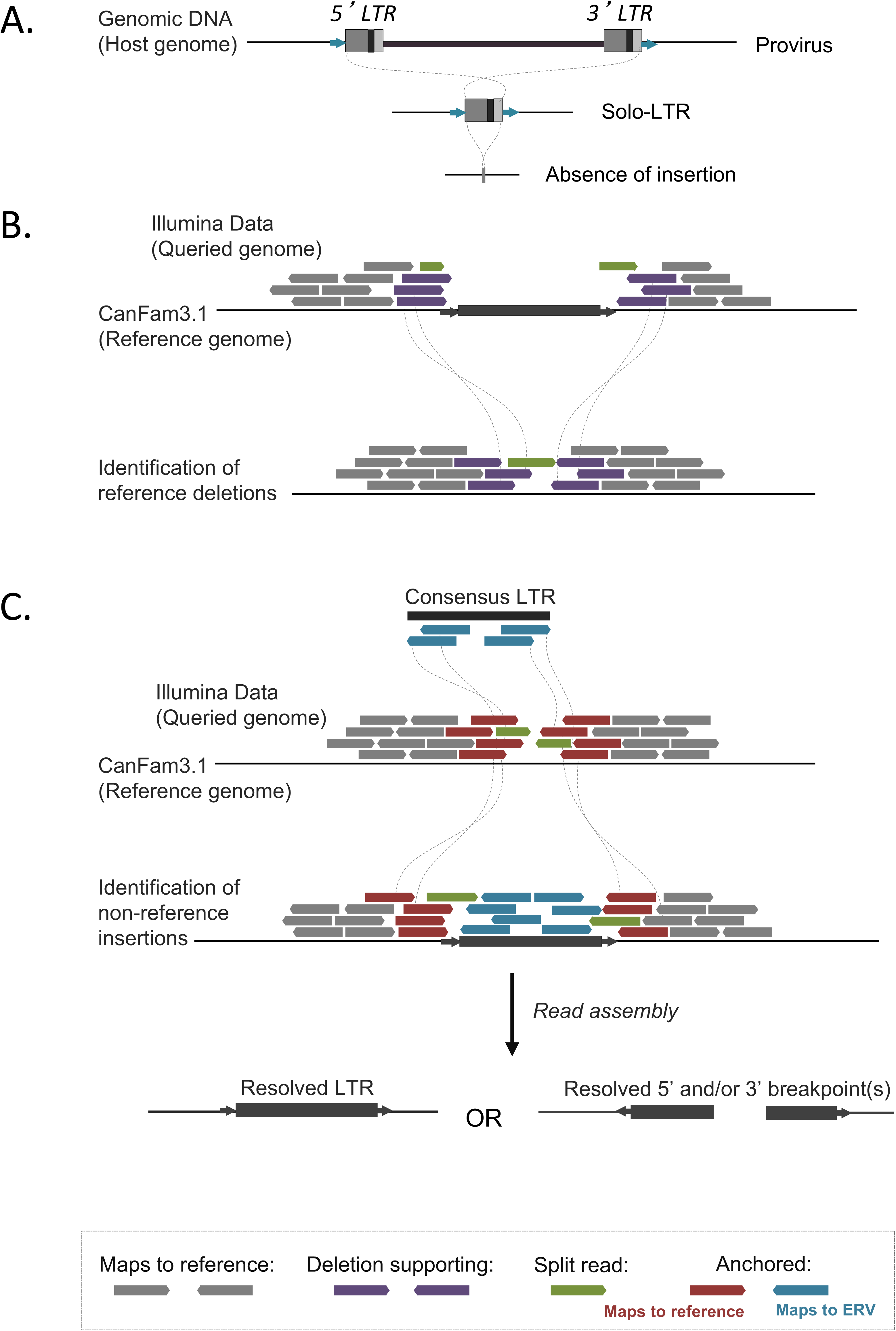
Strategy for detecting insertionally polymorphic ERV variants. (A) ERV allelic presence. Upper: full-length provirus; Mid: solo LTR recombinant; Lower, unoccupied (pre-integration) site. (B) Strategy for detection of reference ERV deletions. Illumina read pairs were mapped to the CanFam3.1 reference, deletion-supporting read pairs and split reads identified using the program Delly (Rausch et al., 2012), and candidate calls then intersected with RepeatMasker outputs considering ‘CFERVF1’ repeats. Deletion calls within a size range corresponding to a solo LTR or provirus were selected for further analysis. (C) Strategy for detection of non-reference ERV insertions. ERV insertion-supporting anchored read pairs were identified from merged Illumina data mapped to the CanFam3.1 reference using the RetroSeq program (Keane et al., 2013). Insertion-supporting read pairs and intersecting split reads were assembled, assemblies for which ‘CfERVF1’ sequence was present were identified by RepeatMasker analysis, and the assembled contigs then re-mapped to the dog CanFam3.1 reference for precise breakpoint identification.

Our second approach utilized aberrantly mapped read-pairs from the same set of 101 genomes to identify CfERV-Fc1(a) copies that are absent from the dog reference genome. We refer to such insertions as ‘non-reference’. These sites were identified using a combined read mapping and *de novo* assembly approach previously used to characterize polymorphic retroelement insertions in humans (Wildschutte et al., 2015, Wildschutte et al., 2016) (Figure 2B; also see Methods). This process identified 58 unique non-reference insertions, all of which derived from ‘CfERVF1’-related elements per RepeatMasker analysis. Twenty-six of the 58 assembled insertion loci were fully resolved as solo LTRs, 30 had non-resolved but linked 5’ and 3’ genome-LTR junctions, and two had one clear assembled 5’ or 3’ LTR junction. Due to the one-sided nature of assembled reads, we note the latter two were excluded from the majority of subsequent analyses (also see Figure S1 and Table S2). The assembled flanking regions and TSDs of each insertion were unique, implying each was the result of an independent germline invasion. Together, our two approaches for discovery resulted in 69 candidate polymorphic CfERV-Fc1(a)-related elements.

#### Validation of allele presence and accuracy of read assembly

We initially surveyed a panel of genomic DNA samples from breed dogs to confirm the polymorphic status of a subset of insertions (Figure 3). We then confirmed the presence of as many of the identified non-reference insertions as possible (34/58 sites) in predicted carriers from the 101 samples, and performed additional screening of each site to discriminate solo LTR and full-length integrants (Table S2). We confirmed a non-reference insertion for each of the 34 sites for which DNA from a predicted carrier was available. A provirus was present at eight of these loci, both insertion alleles were detected at three loci, and a solo LTR was present for the remaining loci. The full nucleotide sequence was obtained for 33 of the 34 insertions, with preference for sequencing placed on the provirus allele when present. The provirus at the final site (chr5:78,331,579) could not be completely spanned due to the presence of highly repetitive sequence within the *gag* gene (∼2,250 bp from the consensus start). We also confirmed the polymorphic nature of the 11 reference CfERV-Fc1(a) insertions predicted to be unfixed, however we did not detect variable insertion states for those sites.

**Figure 3.**
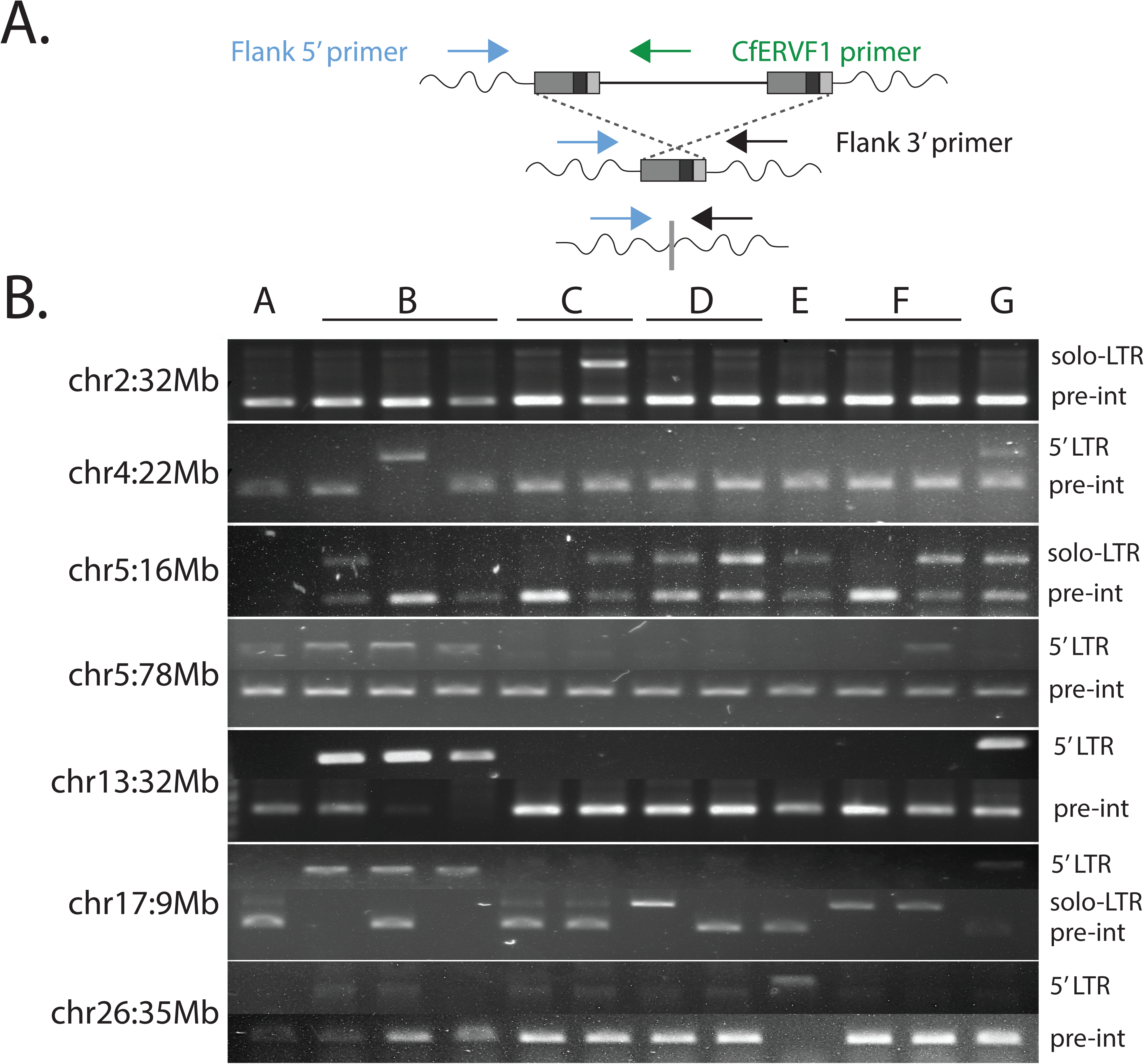
Representative allele screening of polymorphic loci. PCR screens of a subset of non-reference CfERV-Fc1(a) integrants. Validation of insertionally polymorphic sites was performed for seven candidate sites across genomic DNA from a panel of breed dogs. (A) Strategy for primer design and allele detection. Primers were designed to target within 250 bp of the insertion coordinates based on re-mapping of the assembled breakpoints to the CanFam3.1 reference. Two primers sets were used for each locus: one utilized an internal and flanking primer to amplify the 5’ LTR of a full-length element; another set was used for detection of the pre-integration (unoccupied) or solo LTR alleles each locus. (B) Banding patterns supporting the unoccupied, solo LTR, or full-length alleles. The chromosomal location of each integrant is indicated at left; allele presence is indicated at right: (+) insertion presence and detected allele; (-) insertion absence. Samples: A, boxer; B, Labrador retriever; C, golden retriever; D, Springer spaniel; E, standard poodle; F, German shepherd; G, shar-pei.

We assessed the accuracy of read assembly by comparing the assembled alleles to Sanger reads obtained for the validated sites. Due to the inability of the Illumina reads to span a full-length provirus, we were limited to the evaluation of fully assembled solo LTRs. Base substitutions were observed for just two assembled non-reference loci. First, the assembled chr13:17,413,419 solo LTR had a predicted base change between its TSDs that was resolved in Sanger reads; all other validated TSDs were in agreement as 5 bp matches, as is typical of the lineage. Second, the chr16:6,873,790 solo LTR had a single change in the LTR relative to the assembled allele. All other validated loci were in complete agreement with predictions obtained by read assembly of those insertions.

Structural variants between assembled sequences and the reference genome were also observed. For example, the assembled contig at chr33:29,595,068 captured a deletion of a reference SINE insertion 84 bp downstream of the non-reference solo LTR (Figure 4A). Deletion of the reference SINE was also supported by Delly deletion calls using the same Illumina data. Sanger sequencing confirmed a 34 bp deletion in an assembled insertion situated within a TA_(n)_ simple repeat near chr32:7,493,322 (Figure 4B). Finally, an assembled solo LTR that mapped to chr2:32,863,024 contained an apparent 8 bp extension from the canonical CfERVF1 Repbase LTR of its 3’ junction (5’ TTTTAACA 3’). We validated the presence of the additional sequence within matched TSDs flanking the LTR and confirmed its absence from the empty allele (Figure 4C). The extension is similar in sequence to the consensus CfERVF1 LTR (5’ ACTTAACA 3’) and maintains the canonical 3’ CA sequence necessary for proviral integration. These properties support its presence as part of the LTR, possibly generated during reverse transcription or during post-integration sequence exchange.

**Figure 4.**
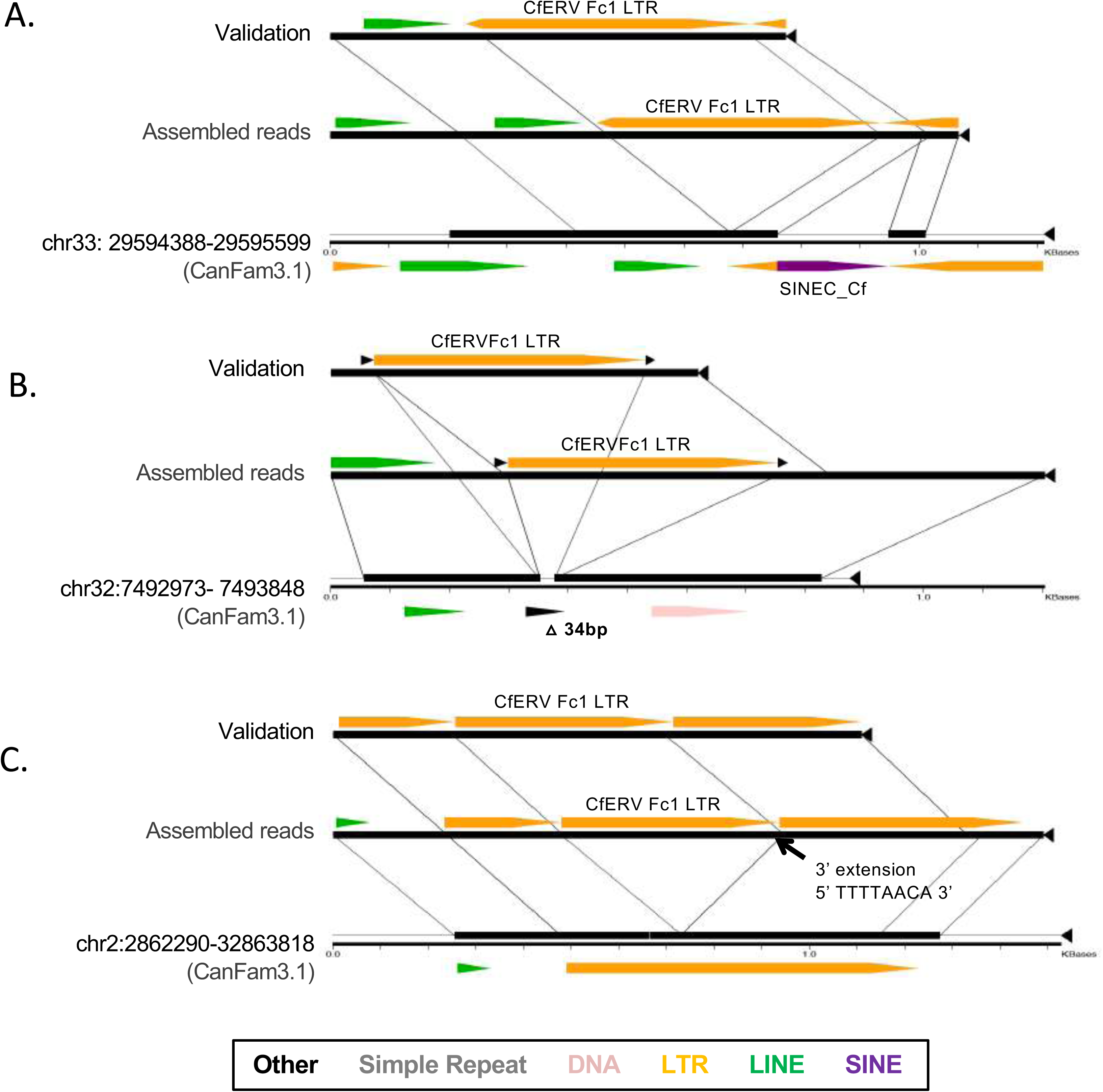
Assessment of assembled non-reference alleles. LTR insertions associated with structural variation as captured in assembled Illumina read data. Local three-way alignments were generated for each assembled locus using the program Miropeats (Parsons, 1995). Each consisted of the LTR allele obtained by read assembly, the validated LTR allele obtained by Sanger sequencing of the locus in one individual, and the empty locus as present within the CanFam3.1 reference. Alignments are shown for three representative LTR assemblies. The allele type is labeled at left in each alignment; lines are used to indicate the breakpoint position of the insertion and shared sequence between alleles. (A) An LTR assembly that includes captured deletion of a bimorphic SINE_Cf insertion present in the CanFam3.1 reference. (B) An assembled LTR associated with a short 34 bp deletion of sequence that is present in the reference. (C) A validated assembly of an LTR that included an 8 bp extension relative to the canonical CfERVF1 repeat.

#### The CfERV-Fc1(a) genomic landscape

In principle, upon integration a provirus contains the necessary regulatory sequences for its own transcription within its LTRs; solo LTR recombinants likewise retain the same regulatory ability. Indeed, ERVs have been shown to affect regulatory functions within the host and some have been exapted for functions in normal mammalian physiology (reviewed in (Jern and Coffin, 2008)). A previous analysis of the then-current CanFam2.0 reference build identified at least five γ-like ERVs within or near genes from proviruses that belonged to a distinct and older non-Fc1(a) sublineage (specifically the ‘CfERV1z’ ERV-P related group, per RepeatMasker) (Martinez Barrio et al., 2011). Given the discovery of numerous novel insertions in our study and the improved annotation of the CanFam3.1 reference assembly, we assessed CfERV-Fc1(a) presence in relation to dog gene models.

Genome-wide insertion patterns were assessed for 58 non-reference and all 107 reference CfERV-Fc1(a) insertions. Of the 165 insertions, 29 (17.6%) were present within the introns of Ensembl gene models while one exonic reference insertion was identified (Table S3). Nine of the genic insertions (30%) were in sense orientation in respect to the gene. Some insertions were also in the vicinity of genes. For example, thirteen additional Fc1 loci were within 5 kb of at least one dog gene model; four of seven insertions situated upstream of the nearest gene were in sense orientation. Another 15 Fc1 loci were within 10 kb of at least one gene, of which seven of ten upstream insertions were in sense orientation with respect to the nearest gene. ERV-related promoter and enhancer involvement has been reported for distances exceeding 50 kb both upstream and downstream of genes (for example, see (Maruggi et al., 2009)). We find that 96 (58.2%) of assessed CfERV-Fc1(a) elements are within 50 kb of a gene model. Compared with randomized placements, CfERV-Fc1(a) insertions are significantly depleted within genes (*p* < 0.001) and within 10 kb of genes (*p* < 0.001). However, no significant difference was observed at the 50 kb distance (Figure S2). Insertions were present on all chromosomes except chr35 and the Y chromosome, which is incomplete and not part of the canonical CanFam3.1 assembly. Individual CfERV-Fc1(a) insertions have been annotated with gene identifiers, gene ontology terms, and distances to nearest gene(s) in Table S3.

### Age and evolutionary relationship of CfERV-Fc1(a) insertions

#### Dating proviral integrants by LTR divergence

Nucleotide divergence between the 5’ and 3’ LTRs of a provirus has been commonly used to estimate the time since endogenization, assuming that ERV sequences evolve neutrally following integration (Johnson and Coffin, 1999, Hughes and Coffin, 2004). Using this dating method, we estimated broad formation times of CfERV-Fc1(a) proviruses that maintained both LTRs. This analysis excluded three truncated reference elements (chr1:48,699,324, chr8:73,924,489, and chrUnAAEX03024336:1) and one non-reference provirus with an internal 291 bp deletion of the 3’ LTR (chr17:9,744,973). The 3’ LTR of the chr33:22,146,581 non-reference insertion contained a 43 bp internal duplication, which we treated as a single change. We applied a host genome-wide dog neutral substitution rate of 1.33×10^−9^ changes per site per year (Botigue et al., 2017), yielding formation times of individual proviruses from 20.49 mya to within 1.64 mya.

These estimates are sensitive to the assumed mutation rate, in addition to the limited number of differences expected between LTRs for the youngest loci. Obtaining age estimates in this manner for the youngest proviruses (as assumed by high 5’ and 3’ LTR identity) is dependent on the time to accrue a single mutation between the LTRs (of ∼457 bp in length). The youngest estimate (1.64 my) is driven by two proviruses whose LTRs differ by a single base change and five proviruses with identical 5’ and 3’ LTRs, although the inter-element LTR haplotype sequence differed between proviruses. Across these five proviruses, LTR identities ranged from 98.5% to 99.4% (average of 98.95%), with a total of five LTR pairs that shared private substitutions. The remaining provirus shared an average identity of 85.45% to the other four. We further identified solo LTRs with sequence identical to one of two respective proviral LTR haplotypes (chr3:82,194,219 and chr4:22,610,555; also see below), suggesting multiple germline invasions from related variants. These data are consistent with insertion of CfERV-Fc1(a) members from multiple exogenous forms in canine ancestors, during which related variants likely infected over a similar timeframe.

#### Prevalence of CfERV-Fc1(a) loci in canids

To more precisely delineate the expansion of the identified CfERV-Fc1(a) members and refine our dating estimates, we surveyed insertion prevalence within an expanded sample set that more fully represent extant members of the *Canidae* family, including the genomes of the dhole (*Cuon alpinus*), dog-like Andean fox (*Lycalopex culpaeus*), red fox (*Vulpes vulpes*), as well as the furthest canid outgroups corresponding to the Island (*Urocyon littorali*) and gray foxes (*U. cinereoargenteus*) (Figure 1). Thus, the analysis provided a broad timeline to reconstruct the evolutionary history of this ERV lineage ranging from host divergences within the last tens of thousands of years (gray wolves) to several millions of years (true foxes).

In total, we *in silico* genotyped 145 insertions (89 reference and 56 non-reference loci) across 332 genomes of canines and wild canids (refer to Methods; Table S4). To more accurately facilitate the identification of putative population-specific CfERV-Fc1(a), and to distinguish possible dog-specific insertions that may have occurred since domestication, wolves with considerable dog ancestry were removed from subsequent analyses (see Methods). Alleles corresponding to reference (*i.e*., CanFam3.1) and alternate loci were recreated based on the sequence flanking each insertion while accounting for TSD presence. We then inferred genotypes by re-mapping Illumina reads that spanned either recreated allele for each site per sample. Reference insertions were deemed suitable for genotyping only if matched TSDs were present with clear 5’ and 3’ LTR junctions. We excluded the two non-reference sites with only a single assembled LTR junction due to uncertainty of both breakpoints (above). To facilitate genotyping of the eight unresolved assemblies with linked 5’ and 3’ LTR junctions, we supplemented the Repbase CfERVF1_LTR consensus sequence over the missing region (lower case in Table S2). As has been discussed in earlier work (Wildschutte et al., 2016), this genotyping approach is limited by the inability of single reads to span the LTR; therefore, the data do not discriminate between the presence of a solo LTR from that of a provirus at a given locus.

Insertion allele frequencies ranged from 0.14% (inferred single insertion allele) to fixed across samples (Figure 5; all raw data is included in Table S5). The rarest insertions were found in gray wolves, the majority of which were also present in at least one village or breed dog (for example, see chr13:16,157,778 and chr15:32,084,977 in Figure 5). All non-reference insertions were variably present in *Canis* species, and only few had read support in outgroup species (*i.e.* foxes, dhole). Notably, there was no evidence for the presence of any loci specific to village or breed dogs. Of outgroup canids, ∼33% (48 of 145) insertions were detected in the Andean fox, and ∼50% (a total of 73) insertions were present in the dhole. The remaining foxes, representing the most distant splits of extant canids, had the lowest prevalence of occupied loci, with just five insertions found in the gray and Island foxes, respectively. However, this is not unexpected since insertions private to these lineages would not be ascertained in our discovery sample set.

**Figure 5.**
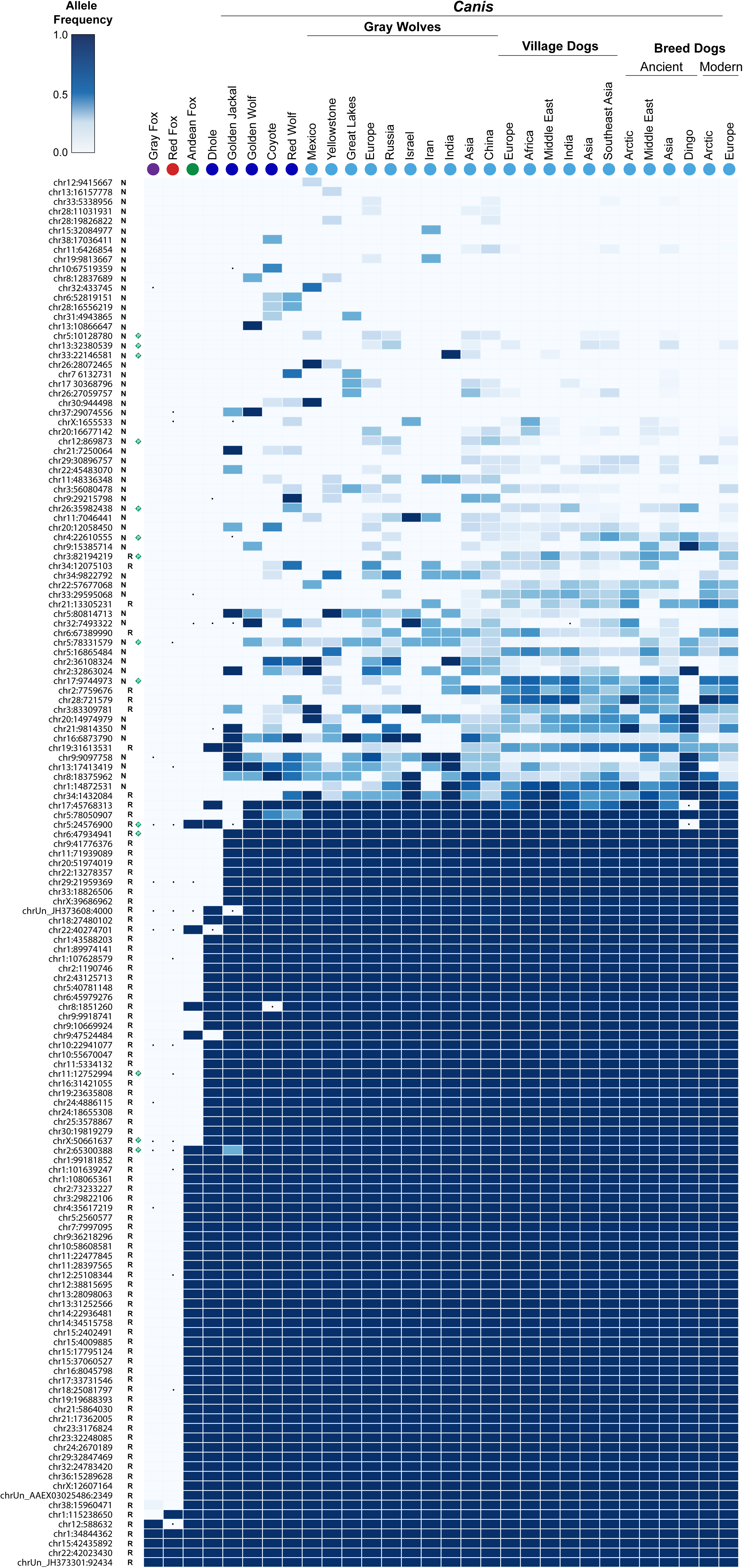
Distribution of CfERV-Fc1(a) insertions in the genomes of modern canids. *In silico* genotyping was performed for 145 LTRs utilizing whole genome data across 347 sequenced canids, which were selected to represent extant members of all major *Canidae* lineages (Figure 1). Sample names are indicated above according to species or sub-population. Samples correspond to the Island and gray foxes (the furthest outgroup species; n=8), red fox (n=1), Andean fox (n=1), dhole (n=1), golden jackal (n=1), golden wolf (n=1), coyote (n=3), red wolf (n=2), and representatives of gray wolf sub-populations (n=33), village dogs (n=111), ancient breed dogs (n=38), and modern breed dogs (n=154). ‘Insertion’ and ‘unoccupied’ alleles were recreated utilizing the CanFam3.1 reference and genotypes were inferred by re-mapping Illumina reads that spanned either recreated allele for each sample. Samples lacking remapped reads across a given site were excluded from genotyping at that site alone (indicated with a ‘.’). Allele frequencies were calculated for each species or sub-population (see Methods) and plotted as a heat map (insertion frequency indicated by color bar at top). The locus identifier for each insertion (left) corresponds to the chromosome and the leftmost insertion breakpoint, irrespective of insertion orientation. Non-reference and reference insertions are indicated by an ‘N’ and ‘R’, respectively. Full-length proviruses are highlighted with a green diamond.

The relative distribution of proviruses was in general agreement with dating via LTR divergence, though some inconsistencies were observed. No proviruses were detected in the fox outgroups (*Urocyon* and *Vulpes*) that have an estimated split time from other Canidae of >8 mya (Kumar et al., 2017), but some were present in the Andean fox (chr2:65,300,388, chr5:24,576,900) and dhole (chrX:50,661,637, chr11:12,752,994). LTR divergence calculations using the inferred dog neutral substitution rate dated these insertions near 20.49, 14.80, 6.65, and 4.94 mya, respectively, suggesting the dating based on LTR divergence may be overestimated. The youngest proviruses were variably present in *Canis* representatives. Of the most recent insertions, two (chr5:10,128,780, chr17:9,744,973) were present in both New and Old World wolves, implying integration prior to the geographic split of this lineage (1.10 mya) (Fan et al., 2016). The remaining proviruses were present in Old World wolves and dogs only. Among these was the chr33:22,146,581 provirus that had an estimated date of formation of 6.58 mya by LTR comparison, consistent with skewed dating of the site. Altogether, the data are consistent with CfERV-Fc1(a) endogenization in the ancestors of all modern canids followed by numerous invasions leading to a relatively recent burst of activity in the wolf and dog lineage of *Canis*.

#### Evolution of the CfERV-Fc1(a) lineage in Canidae

LTR sequences are useful in a phylogenetic analysis for exploring the evolutionary patterns of circulating variants prior to endogenization, as well as following integration within the host. To infer the evolutionary history leading to CfERV-Fc1(a) presence in modern canids, we constructed an LTR tree using as many loci as possible (from 19 proviral elements and 142 solo-LTRs) (Figure 6; Table S6).

**Figure 6.**
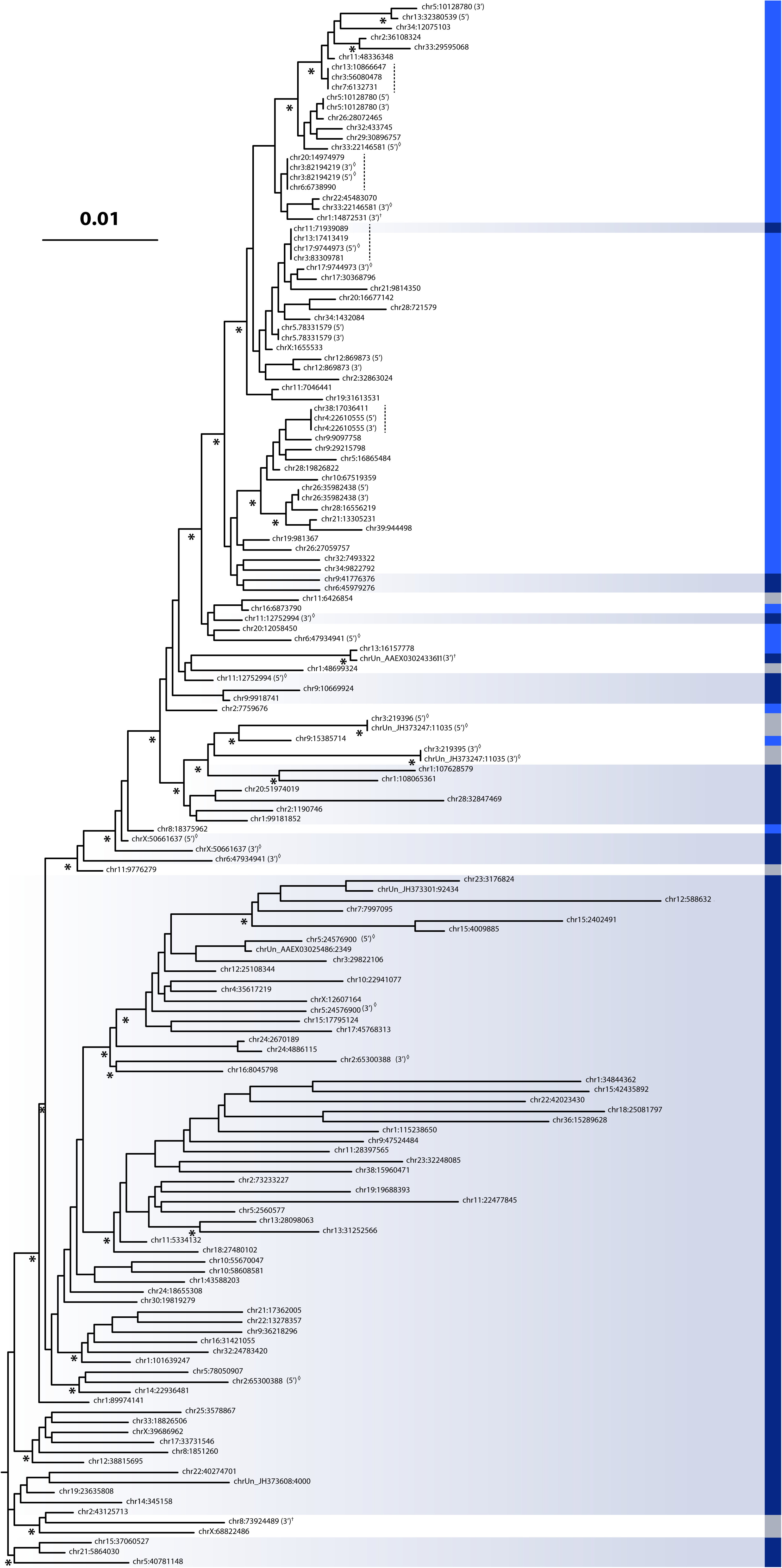
Evolutionary history of the CfERV-Fc1(a) lineage in canids. An approximately-maximum-likelihood phylogeny was reconstructed from an alignment of 157 ERV-Fc LTR sequences. The tree has been midpoint-rooted for display purposes. Asterisks below nodes indicate local support values > 70%. Each insertion is denoted corresponding to chromosomal position relative to CanFam3.1 coordinates. A color bar is shown at the right to denote element presence as fixed among *Canis* (dark blue), insertionally polymorphic (light blue), or not genotyped (gray). Elements identified as fixed among *Canis* insertionally polymorphic have been further highlighted by blue shading. LTRs belonging to proviruses are indicated along with the chromosomal position with a (5’) or (3’) as appropriate. Clusters of identical LTR haplotypes are indicated with a vertical dashed line. Mis-matched pairs of proviral LTRs are indicated by a diamond. LTRs from proviruses lacking cognate LTR pairs (*i.e*., due to truncation of the element) are indicated with a cross. The scale bar shown represents the evolutionary distance in substitutions per site.

In broadly comparing LTR placement to our inferred species presence (Figure 6), the longer-branched clusters contained the few ancestral loci present in the outgroups (gray and red foxes) and those that were mostly fixed among the other surveyed species. However, at least two non-reference LTRs and other unfixed insertions were also in these clades, suggesting their more recent formation from related variants therein. One provirus was present within the most basal clade, and four (including the duplicated locus) were present within intermediate clades. We observed a major lineage (upper portion of tree) that included the majority of recent integrants. This lineage gave rise to the greatest number of polymorphic insertions, including a derived clade of insertions that appears to be *Canis*-specific, with some sites restricted to one or two sub-populations. This lineage also contains the majority of proviral LTRs (15 of 19 included in the analysis), most possessing intact *pol* and/or *env* genes. The youngest proviral integrants, as inferred from high LTR identities and prevalence among sampled genomes, tend to be on short branches within derived clusters that contain the majority of unfixed loci, likely reflecting their source from a relatively recent burst of activity in *Canis* ancestors.

Within the germline, the highest occurrence of recombination resulting in a solo LTR takes place between identical LTRs (Belshaw et al., 2007, Stankiewicz and Lupski, 2002), implying the LTR sequence itself is preserved in the solo form. Under this assumption, the presence of identical solo LTR haplotypes should, in principle, indicate their origin from a common ancestral source. We identified four such LTR haplotypes within the *Canis*-specific clades, including loci in co-clusters with one of two proviruses (chr3:82,194,219 and chr4:22,610,555), therefore bounding the inferred age of these insertions to within the last 1.64 mya (dashed lines in Figure 6). Between the four identical clusters, the LTR haplotypes shared nucleotide identity ranging from 99.3% (three substitutions from a consensus of the four clusters) to 99.7% (one substitution), suggesting their origin from related variants over a common timeframe. We modified our dating method to obtain an estimated time of formation across each cluster by considering the total concatenated LTR length per cluster, as has been similarly employed elsewhere (Ishida et al., 2015). This approach placed tentative formation times of the youngest insertions from a common variant 547,220 years ago (no change over 1,374 bp, or 3 LTRs) and 410,415 years ago (no change over 1,832 bp, or 4 LTRs). Comparison to the inferred prevalence of each cluster indicates the most recent of these insertions arose in Old World wolves, consistent with this timeframe.

Since proviral LTRs begin as an identical pair, aberrant placement in a tree and/or the presence of mismatched TSDs implies involvement of the locus in post-insertion conversion or rearrangement (Hughes and Coffin, 2005). LTRs from the youngest proviruses tended to pair on sister branches. An exception includes the LTRs of the chr33:22,146,581 provirus, whose mispairing is consistent with conversion of at least one of its LTRs, possibly from the chr1:48,699,324 provirus or a similar variant (see above). There were six instances of aberrant LTR placement for the remaining eight CfERV-Fc1(a) proviruses that had both LTRs present (labeled in Figure 6), suggesting putative post-insertion conversion and contributing to inflated age estimates based on LTR divergence. The TSD repeats of individual proviruses had matched 5 bp repeats in all cases, suggesting none of the elements have seeded inter-element chromosomal rearrangements. With exception of three instances of reference solo LTRs that each had a base change between its flanking repeats, the TSDs for all other solo LTRs were also intact.

### CfERV-Fc1(a) structure and biology

#### Characterization of the inferred CfERV-Fc1(a) ancestor

As an endogenous element, an ERV may retain close resemblance to its exogenous source over long periods of time with recent integrants assumed to possess a greater potential to retain the properties of the infectious progenitor. We combined the eight non-reference proviruses with the eleven reference insertions to generate an updated consensus (referred to here as CfERV-Fc1(a)_CON_) as an inferred common ancestor of the CfERV-Fc1(a) sublineage. A detailed annotation of the updated consensus is provided in Figure S3 and summarized as follows.

Consistent with the analysis of Caniform ERV-Fc1 consensus proviruses (Diehl et al., 2016), CfERV-Fc1(a)_CON_ shows an internal segment of uninterrupted ERV-Fc related ORFs for *gag* (∼1.67 kb in length) and *pol* (∼3.54 kb; in-frame with *gag*, beginning directly after the *gag* stop codon, as is typical of C-type gammaretroviral organization). The CfERV-Fc1(a)_CON_ *gag* product was predicted to contain intact structural regions and functional motifs therein for matrix (including the PPPY late domain involved in particle release and the N-terminal glycine site of myristoylation that facilitates Gag-cell membrane association), capsid, and nucleocapsid domains (including the RNA binding zinc-binding finger CCHC-type domains). Likewise, the Fc1(a)_CON_ *pol* ORF was predicted to encode a product with conserved motifs for protease, reverse transcriptase (the LPQG and YVDD motifs in the RT active center), Rnase H (the catalytic DEDD center of RNA hydrolysis), and integrase (the DDX_35_E protease resistant core and N-terminal HHCC DNA binding motif). An *env* reading frame (absent from the Repbase CfERVF1 consensus) was also resolved in the updated consensus. The ERV-W like Fc1_CON_ *env* ORF (∼1.73 kb) was present within an alternate ORF overlapping the 3’ end of *pol*. Its predicted product included the RRKR furin cleavage site of SU and TM, the CWIC (SU) and CX_6_CC (TM) motifs involved in SU-TM interactions, and a putative RD114-and-D-type (RDR) receptor binding motif (Sinha and Johnson, 2017). A hydrophobicity plot generated for the translated sequence identified segments for a predicted fusion peptide, membrane-anchoring TM region, and immunosuppressive domain (ISD) (Cianciolo et al., 1985). Putative major splice donor (base 576 within the 5’UTR; 0.67 confidence) and acceptor sites (base 5,216 within *pol*; 0.85 confidence) were identified that would be predicted for the generation of *env* mRNA (see Figure S3 and the accompanying legend). The CfERV-Fc1(a)_CON_ element possessed identical LTRs, a tRNA^Phe^ binding site for priming reverse transcription (GAA anticodon; bases 464 to 480), and the canonical 5’-TG…CA-3’ terminal sequences required for integration (Boeke and Stoye, 1997).

#### Properties of individual CfERV-Fc1(a) proviruses

We assessed the properties of individual full-length elements for signatures of putative function (summarized in Figure 7). With the exception of the *gag* gene, we identified intact ORFs in several reference copies and most of our non-reference sequenced proviruses. A reading frame for the *pol* gene was present in six proviruses; of these, all contained apparent RT, RnaseH, and integrase domains without any changes that would obviously be alter function. Likewise, an *env* ORF was present among seven proviruses, of which all but one contained the above mentioned functional domains (the SU-TM cleavage site is disrupted in the chr5:10,128,780 provirus: RRKA). Comparison of the rate of nonsynonymous (d_N_) to synonymous (d_S_) nucleotide substitutions for the seven intact *env* reading frames revealed an average d_N_/d_S_ ratio of 0.525, indicating moderate purifying selection (*p* = 0.02, Nei-Gojobori method). The hydrophobicity plot of each *env* ORF was in agreement with that of the CfERVFc1_CON_ provirus, with predicted segments for a fusion peptide, TM region, and ISD. Comparison to the *pol* and *env* translated products that would be predicted from the CfERVFc1_CON_ inferred the individual proviruses shared 98.4% to 99.3% (Pol) and 98% to 99.6% (Env) amino acid identity, respectively, and each was distinct from the inferred consensus.

**Figure 7.**
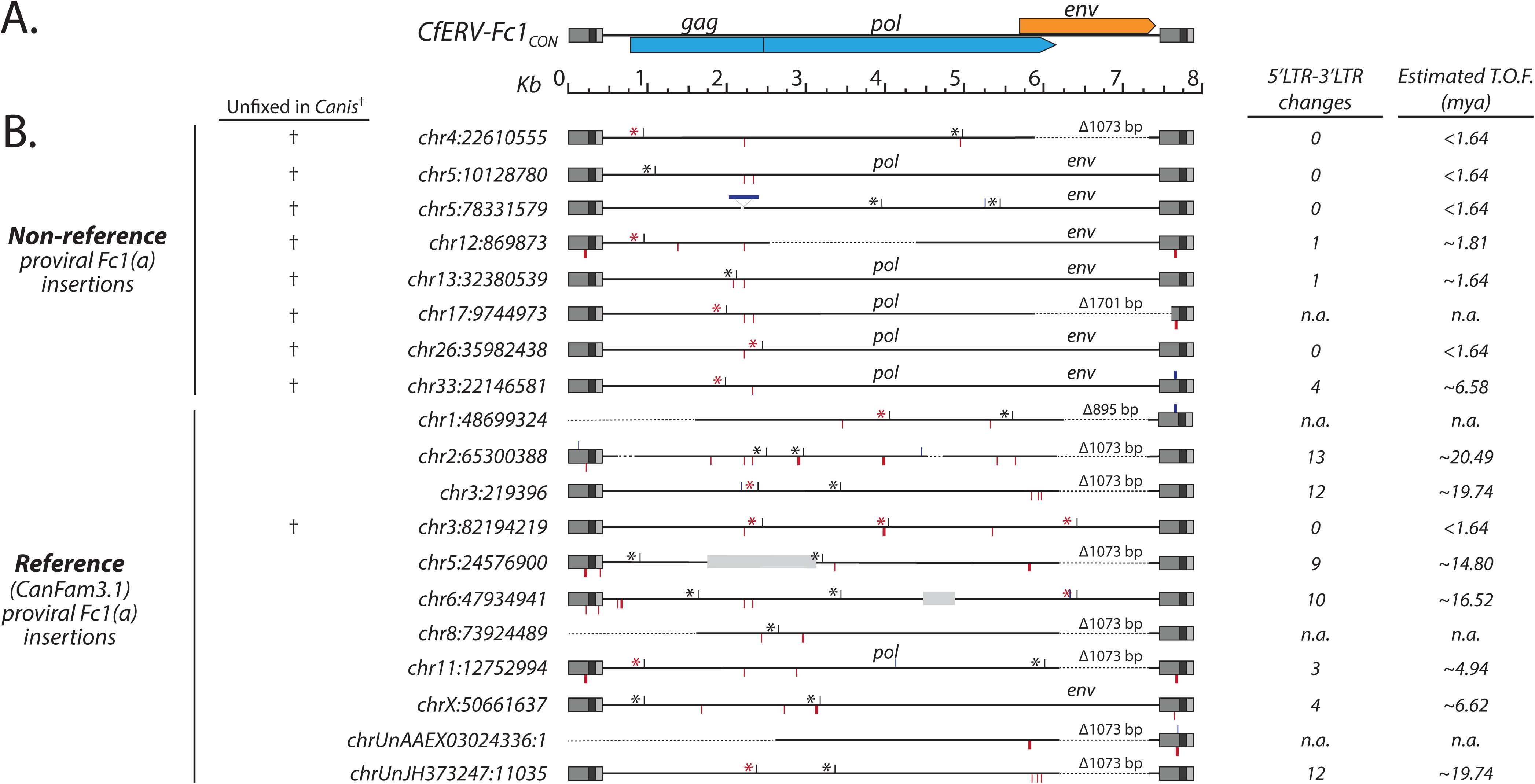
Structural features of CfERV-Fc1(a) proviruses. (A) Representation of the CfERV-Fc1(a)_CON_ provirus. Viral gene reading frames for ERV-Fc related *gag* and *pol* are shown in blue; the ERV-W related *env* is shown in orange. Color usage is consistent with that of (Diehl et al., 2016). LTRs are colored in gray: U3 is in medium tone; R is dark; U5 is light. The provirus and open reading frames are shown to scale. (B) Structural features of non-reference and reference proviruses. When present, open reading frames are indicated above the appropriate element. Insertions and deletions >3 bases are depicted with blue and red flags, respectively. The *env*_Δ1073_ deletion is labeled and indicated by a dashed line, as are other truncated or deleted element features. Reference gaps present within are shown in light gray boxes to scale. Stop codons are indicated with a black or red asterisk, where red is used to specify premature stops common to two or more proviruses. Crosses at the left indicate proviruses that are unfixed among *Canis* samples. The number of substitutions between LTRs is shown at right with the corresponding calculated age as inferred based on the neutral substitution rate of 1.33×10^−9^ changes per site per year (Botigue et al., 2017).

No complete *gag* reading frame was observed in any provirus. Particularly when compared to *pol* and *env*, the *gag* gene had incurred a number of inactivating mutations, including several shared frameshifts leading to premature stops. The longest *gag* reading frames (chr3:82,194,219 and chr26:35,982,438) both possessed a premature stop within the first zinc finger domain of the nucleocapsid; of note the terminal *gag* frameshift was the only obvious inactivation of any gene in the latter chr26:35,982,438 provirus. This domain has roles in the encapsidation of viral genomic RNAs via recognition of a particular packaging signal sequence (Ali et al., 2016). Thus, absence of both zinc finger domains and the N-terminal myristoylation site should interfere with canonical Gag functions, regardless of the presence of intact matrix and capsid domains. Excluding the frameshift leading to the abortive stop in those proviruses, the translated Gag would have respectively shared 97.8% and 98% amino acid identity to the CfERVFc1_CON_ Gag. Though none of the identified CfERV-Fc1(a) proviruses have retained complete reading frames for all genes, this finding does not exclude the possibility that rare intact proviruses remain to be identified, or that a putative infectious variant could be generated via recombination of co-packaged RNAs.

The majority of the CfERV-Fc1(a) proviruses could be assigned to one of two proposed subgroups based on the presence of a common deletion within the *env* gene (Figure 7). The deletion spans a 1,073 bp region of *env* (we refer to the segment as *env*_Δ1073_), removing the internal majority portions of SU and TM (also refer to Figure S3; including the putative receptor binding domain, motifs involved in SU-TM interactions, and transmembrane domain). Eight proviruses possessed the *env*_Δ1073_ deletion, including the duplicated locus. The frameshift for three of those eight proviruses would result in a product of ∼204 amino acids in size (chr2:65,300,388, chr4:22,610,555, and chr6:47,934,941), though the significance of such a product is unclear. The prevalence of the *env*_Δ1073_ deletion was skewed toward proviruses that harbored multiple inactivating mutations, while only one possessed a retained ORF (chr11:12,752,994, *pol*), consistent with the older status of most of these loci. Additionally, the *env*_Δ1073_ deletion was present in the oldest proviruses and inferred to have arisen at least prior to the split of the dog-like foxes (see chr2:65,300,387 in Figure 5), suggesting its formation early in CfERV-Fc1(a) evolution (at least 8.7 mya; Figure 1). However, three proviruses with the deletion could not be genotyped due to the absence of clear LTR-genome junctions or due to encompassing duplication, making it possible that the allele predates the Andean fox split, as would be consistent with their placement within the tree (for example, see chr8:73,924,489; Figure 6). The *env*_Δ1073_ deletion was not monophyletic in gene or LTR-based phylogenies, as would be expected if proviruses carrying the allele arose from a ‘master’ source element (Clough et al., 1996, Nascimento and Rodrigo, 2016). Examination of the regions directly flanking the deletion did not reveal common base changes shared among members with the allele. Our data are also not consistent with its transfer to existing proviruses through gene conversion, which should display shared base changes between all elements with the deletion. Therefore, we propose the *env*_Δ1073_ allele spread via template-switching of co-packaged *env*_Δ1073_ RNAs. Any of the above scenarios would result in the spread of an otherwise defective *env* gene. In contrast, all but two of the most recently integrated proviruses contained an uninterrupted *env* reading frame. In addition to the *env*_Δ1073_ deletion, unique *env* deletions were present in two other elements. Specifically, a 1,702 bp deletion which removed all but the first 450 bp of *env* and 291 bp of the chr17:9,744,973 3’ LTR, as well as the 5’ truncated provirus at chr1:148,699,324 with an 896 bp deletion situated within the common *env*_Δ1073_ deletion.

#### CfERV-Fc1(a) proliferation in canine ancestors

Nucleotide signatures within ERVs may be used to infer the mode(s) of proliferation, of which several routes have been described. One such mechanism, *trans* complementation, involves the co-packaging and spread of transcribed viral RNA genomes by functional viral proteins, supplied by a virus within the same cell (either exogenous or endogenous), thereby prolonging the apparent activity of the ERV lineage. As a result, RNAs from otherwise defective proviruses may be spread in cases where the ERV retains intact structures for transcription by host cell machinery and RNA packaging (Boeke and Stoye, 1997). Molecular signatures of *trans* complementation may be interpreted from the presence of inherited changes among multiple elements, particularly ones that would render a provirus defective (Belshaw et al., 2004, Belshaw et al., 2005).

We observed evidence for the mobilization of CfERV-Fc1(a) copies via complementation. For example, examination of the proviral gene regions revealed inherited frameshift-causing indels and common premature stops that were variably present among the majority of elements (a total of 12 of the 19 proviruses; also see Figure 7). At least three distinct frameshifts leading to a stop within *gag* were shared over several elements (from the Fc1_CON_ start, bp 882: chr4:22,610,555, chr11:12,752,994, chr12:869,873; bp 1,911: chr17:9,744,973, chr33:22,146,581; and bp 2,203: chr3:82,194,219, chr26:35,982,438, and the duplicated chr3:219,396 and chrUn_JH373247:11,035 insertions). Proviruses also shared unique deletions leading to abortive stops within *pol* (near Fc1_CON_ bp 3,988: chr1:48,699,324, and chr3:82,194,219). In addition to the common *env*_Δ1073_ frameshift deletion, putative in-frame *pol* deletions were also present (Fc1_CON_ bp 5,263 Δ3 bp: chr3:82,194,219; chrUn_AAEX03024336:1; bp 5,705 Δ27 bp: chr5:24,576,900, chrUn_AAEX03024336:1). Two proviruses contained a shared stop within *env* (Fc1_CON_ bp 6,240: chr3:82,194,219, chr6:47,934,941). The provirus on chromosome 3 possessed a total of four of the above changes differentially shared with other proviruses in *gag pol*, and *env*; these were the only defective changes present within the element. While successive conversion events of the provirus from existing loci cannot be ruled out, this provirus appears to be a comparatively young element (only found in Old World wolves and dogs), which more likely suggests formation of the element via multiple intermediate variants. No other provirus contained multiple common indels.

We did not find evidence for expansion of the lineage having proliferated via retrotransposition in *cis*, during which new insertions are generated in an intracellular process akin to the retrotransposition of long interspersed elements (Ostertag and Kazazian, 2001). Such post-insertion expansion is typically accompanied by a loss of the viral *env* gene, particularly within recently mobilized insertions (as interpreted, for example, by the derived phylogenetic placement), whereas *gag* and *pol* are retained. Our data suggest this scenario is unlikely given the absence of a functional *gag* gene and presence of a conserved *env* ORF in several elements, particularly young ones. In this regard, *cis* retrotransposition tends to facilitate rapid *env*-less copy expansion and therefore tends to occur among derived copies of a given lineage (Magiorkinis et al., 2012), and our data suggest the opposite regarding older (loss of *env*) and younger (*env* present) CfERV-Fc1(a) proviruses.

## Discussion

Mammalian genomes are littered with the remnants of retroviruses, the vast majority of which are fixed among species and present as obviously defective copies. However, the genomes of several species harbor some demonstrably ancient ERVs whose lineages contain relatively intact loci and are sometimes polymorphic, despite millions of years since integration. Such ERVs have the potential to exert expression (of either proviral-derived or host-derived products by donation of an LTR) that may affect the host, especially for intact copies or those within new genomic contexts. In particular, ERV expression from relatively recent integrants has been linked to disease (reviewed in (Jern and Coffin, 2008, Mager and Stoye, 2015)). However, there is also growing evidence that many fixed loci have been functionally co-opted by the host or play a role in host gene regulation (reviewed in (Frank and Feschotte, 2017)). To begin to discriminate the relationship of individual elements in the context of the host necessitates a population-level investigation of the breadth of ERV abundance and prevalence within a species. Illustrating both bursts of activity and putative extinction, our findings present a comprehensive assessment of the evolutionary history of a single retroviral lineage through the genomic surveys of nine globally distributed canid species, some represented by multiple subpopulations.

Relative to other animal models, ERV-host relationships within the dog have been understudied. Until now, reports of canine ERVs have been from analysis of a single genome assembly or limited screening of reference loci (Martinez Barrio et al., 2011, Jo et al., 2012, Tarlinton et al., 2013). To further investigate a subset of apparent recent germline integrants (Martinez Barrio et al., 2011) we surveyed the level of polymorphism and possible mechanisms of spread of the γ-like ERV-Fc1(a) lineage across a diverse set of canid species. Our exhaustive analysis of CfERV-Fc1(a) loci is the first population-level characterization of a recently active ERV group in canids. From analysis of Illumina WGS from 101 representatives from the canid genus *Canis*, we uncovered numerous insertionally polymorphic sites, corresponding to both reference loci and insertions that are missing from the dog genome assembly, and performed a comparative genotyping approach utilizing additional extant *Canidae* members to provide a broad evolutionary history of this ERV lineage. We identified eight non-reference proviruses that contain ORFs, display high LTR identities, and have derived placements within a representative phylogeny, which are all characteristics of relatively young elements.

Insertions were located within dog gene models, although permutations indicated that CfERV-Fc1(a) insertions are significantly depleted within and near genes (Figure S2). Given their placement in previously unoccupied genomic locales, the presence of such insertions raises the possibility of biological effects. For example, two intronic LTRs were fixed in all canids: one within *AIG1*, a transmembrane hydrolase involved in lipid metabolism (Parsons et al., 2016); the other in the diffuse panbronchiolitis region *DPCR1* of the dog major histocompatibility complex 1 (Yan et al., 2018). Other intronic insertions were fixed in samples following the splits of the true and dog-like foxes. These included genes with homologs involved in tumor suppression (*OPCML*), cell growth regulation (*CDKL3*), DNA repair (*FANCL*), and innate immunity (*TMED7*-*TICAM2*). An exonic *Canis*-specific solo LTR was located at chr1:107,628,579 within the 3’ UTR of *BCAT2*, an essential gene in metabolizing mitochondrial branched-chain amino acids. In humans, altered expression of *BCAT2* is implicated in tumor growth and nucleotide biosynthesis in some forms of pancreatic cancer (Mayers et al., 2016, Dey et al., 2017, Ananieva and Wilkinson, 2018). The same LTR is situated ∼550 bp upstream of *FUT2*, a fucosyltransferase involved ABH blood group antigen biosynthesis in mucosal secretions (de Mattos, 2016, Ferrer-Admetlla et al., 2009). *FUT2* variants affect secretion status and have been implicated in intestinal microbiota composition (Le Pendu et al., 2006), viral resistance (Thorven et al., 2005), and slowed progression of HIV (Kindberg et al., 2006). Other insertions were upstream and downstream of gene vicinities, again raising the possibility of host effects (also refer to Table S3). Though connections between LTR presence and physiology are yet to be determined, these findings will inform future investigations into the potential effect of CfERVs on host biology.

CfERV-Fc1(a) integrants endogenized canid ancestors over a period of several millions of years (Figure 8B-E). This activity included bouts of infectious activity/mobilization inferred from the last 20.4 my to the most recent integrants formed within 1.6 mya, the latter of which are only present in *Canis* sub-populations. The mutation rate we used to obtain these estimated timeframes (1.33×10^−9^ changes per site per year (Botigue et al., 2017)) coincides with those from two other ancient genome analyses, which utilized ancient DNA to calibrate wolf and dog mutation rates (Skoglund et al., 2015, Frantz et al., 2016). However, our rate is substantially slower than those used previously to date reference CfERV-Fc1(a) members including 2.2×10^−9^ (as an “average” mammalian neutral substitution rate) (Martinez Barrio et al., 2011) and the faster rate of 4.5×10^−9^ (as has been reported for the mouse) (Diehl et al., 2016). Applying those substitution rates to our data would infer much younger integration times of 11.85 mya to <0.91 mya and 6.1 mya to <0.48 mya, respectively. We note the precision in ERV-Fc1(a) age estimations using this method is subject to the accuracy of the inferred background mutation rate, but may be skewed due to the presence of post-insertion sequence exchange between LTRs. As the latter cannot be conclusively ruled out, we interpret our estimations as broad formation times only.

**Figure 8.**
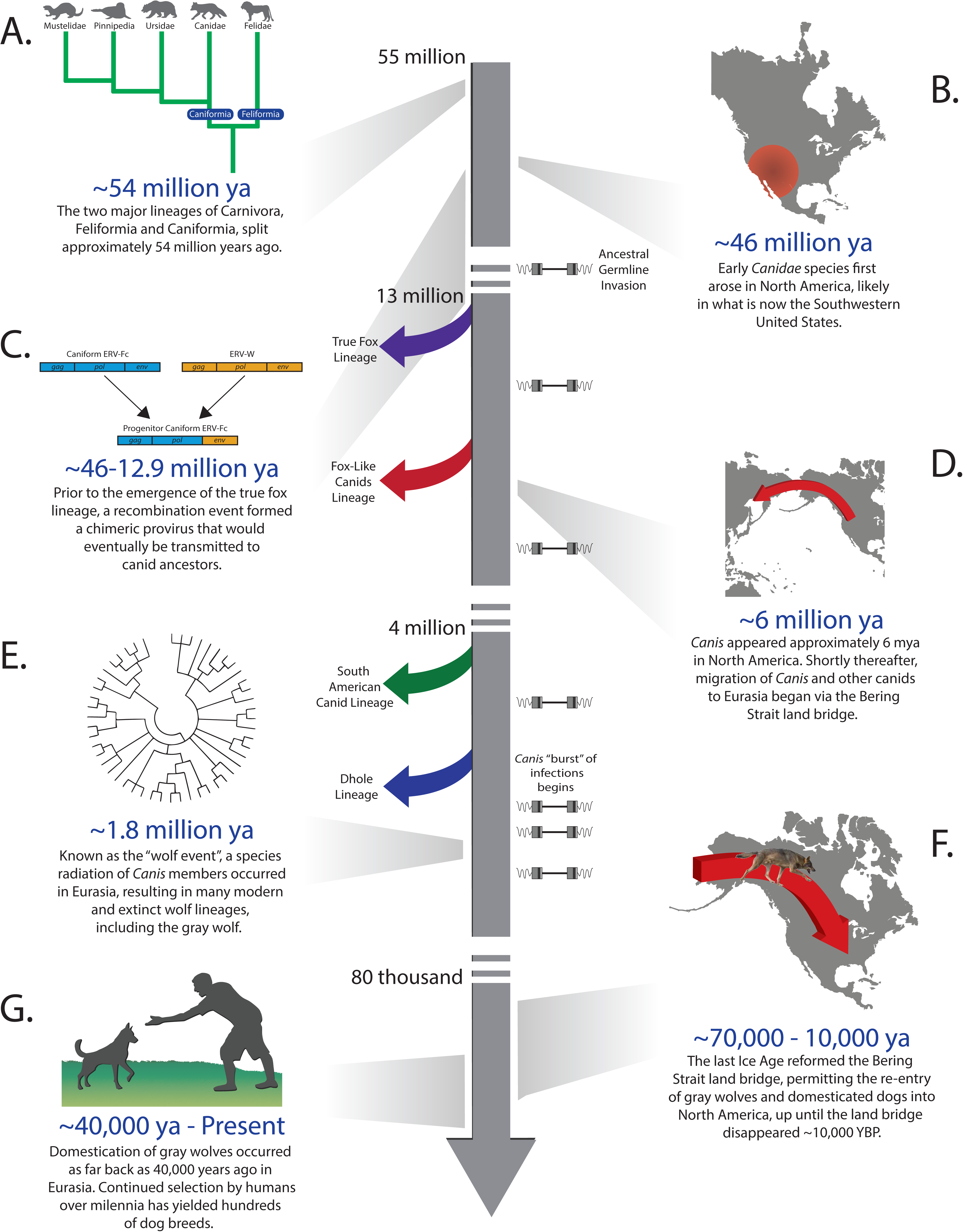
History of CfERV-Fc1(a) germline invasion in the Canidae. A timeline of major events in canid or CfERV-Fc1(a) evolutionary history relative to estimated insertion events. At the approximate time point, branching events of the major canid lineages are indicated by arrows along the timeline with colors matching Figure 1. Indicated by proviruses to the right of the timeline are estimated insertion times based on genotyping data from Figure 5. (A) Based on its presence in all canids, the recombination event that formed the provirus (B), which infected canid ancestors occurred sometime between the split of the major Caniform lineages (A) and the origins of canids in North America (C). Following the migration to Eurasia (D), a major species radiation occurred in the wolf-like canid lineage (E). Finally, the comparatively recent re-introduction of gray wolves in North America reflects the split between the Old and New World wolves (F), which likely partially coincided with the domestication of Old World Wolves (G). Estimated timings for events A-C are supported by (Kumar et al., 2017), D-E by (Wang and Tedford, 2008), F by (Koblmüller et al., 2016), and G by (Botigue et al., 2017).

Due to their complete absence of LTR divergence, the youngest CfERV-Fc1(a) ages are bounded to the estimate of 1.64 my, using the dog substitution rate. Therefore, we employed an alternative approach that makes use of LTRs that shared haplotypes (Ishida et al., 2015) to narrow the age estimations to ∼547,220 and 410,415 years, again, as inferred from the time estimated to accrue one mutation across multiple identical LTRs (respectively across three and four LTRs per haplotype). For comparison, applying the average mammalian and mouse substitution rates to the same data would place either event respectively at 303,251 and 161,734 years ago (no change over three LTRs) and 227,438 and 121,300 years ago (no change over four LTRs). Both estimates are consistent with CfERV-Fc1(a) circulation after the estimated emergence of the gray wolf species 1.1 mya and pre-dating the split of the New and Old World gray wolves (Fan et al., 2016) (Figure 8F). The branching patterns observed within our LTR phylogeny are consistent with these findings, implying bursts of replication from closely related variants now recorded in clusters of LTR haplotypes. In this regard, our findings suggest bouts of infection from multiple circulating viruses over a relatively short evolutionary time period.

CfERV-Fc1(a) activity coincided with major speciation events in canine evolution (Figure 8B-E). Taking into consideration the above approaches for age estimations, we refined the dating of endogenization events by integrating inferred ages with that of orthologous presence/absence patterns across numerous canid lineages, many of which are recently diverged clades. The analysis served two purposes. First, we made use of the tenet that ERV integration is permanent and the likelihood of two independent integration events at the same locus is negligible. In this way, the presence of an ERV insertion that is shared between individuals or species supports its origin in a common ancestor. Therefore, integration prior to or following the split of two or more species is supported by virtue of insertion presence/absence of occupied loci across those species. Second, the analysis allowed us to infer insertion genotypes across highly diverse canid representatives, thus providing the means to gauge the collective patterns of individual CfERV-Fc1(a) loci among contemporary animals to infer putative sub-population or species-specific integrants.

Comparisons of the approximated insertion dates discussed above in combination with estimated species split times would place the earliest CfERV-Fc1(a) germline invasions prior to or near the estimated divergence of the *Canidae* from now extinct ancestors (14.15 mya) (Kumar et al., 2017), followed by invasions after the split of the true fox (12.9 mya) (Kumar et al., 2017) and fox-like canid lineages (8.7 mya) (Koepfli et al., 2015). Subsequent insertions also occurred prior to the split of the South American canid and wolf lineages (3.97 mya) (Koepfli et al., 2015). According to this timeframe, and consistent with the detection of some young proviral insertions private to gray wolves and dogs alone (Figure 5), the most recent invasions would have occurred around the time of the branching event that gave rise to gray wolves (1.10 mya) (Koepfli et al., 2015). Based on the lack of observed dog-specific loci, our data suggests that CfERV-Fc1(a) replication ceased in wolf ancestors prior to domestication, which is estimated to have begun around 40 kya (Botigue et al., 2017) (Figure 8G), but does not rule out continued activity. Analysis of additional genomes, particularly from gray wolves, should clarify the presence of such variants in future analysis.

ERV-Fc1(a) activity included the spread of defective recombinants. Our comparative analysis of nucleotide differences shared among the proviruses supports a scenario in which CfERV-Fc1(a) members proliferated in canine ancestors via complementation. Patterns of discreet, shared changes among distinct elements in all viral genes were observed (*i.e*., premature stops and common base changes, indels, in addition to the *env*_Δ1073_ segment; Figure 7), consistent with the spread of mutations present from existing Fc1(a) copies, probably via co-packaging of the defective viral genomes. Of the 19 proviruses analyzed in full, the majority displayed shared discreet stops or the *env*_Δ1073_ deletion, in addition to in-frame indels. This pattern is consistent with the hypothesis that degradation of ERV genomes, particularly involving the loss of *env*, offers an evolutionary benefit to the host by preventing the potential horizontal spread of infectious viruses between individuals, as has been suggested (Magiorkinis et al., 2012, Lober et al., 2018). The presence of intact *env* genes, and sequence signatures of selective pressure retained within those *env* reading frames, suggests involvement of Fc1(a) *env* leading to the putative formation of recombinant proviruses, rather than having been intracellularly retrotransposed (in *cis*) that would not require a functional envelope. Altogether such patterns of reinfection may have predominantly occurred within a given individual, as none of these mechanisms explicitly requires (but does not rule out) spread to other individuals within the population; indeed concurrent reinfection of a single individual may also lead to unique proviruses later transmitted to offspring (Young et al., 2012). Indeed, several retroviruses, including HIV, have been shown to be capable of co-packaging RNA from other retroviruses, even ones with low sequence homology (Ali et al., 2016). These findings suggest that complementation was a predominant form of proliferation for the observed CfERV-Fc1(a) loci. In theory, a functional provirus could arise in a spontaneous recombinant, raising the possibility of bursts of amplification to come. Indeed, all viral genes in our consensus appear to be intact, illustrative that few changes would be required to generate a putatively infectious virus.

Patterns of shared sequence changes, such as premature stops and in-frame shifts, indicate that the oldest inherited change involved an in-frame shift in the *pol* gene (from the Fc1_CON_ start, bp 5705 Δ27 bp). Aside from the *env*_Δ1073_ deletion, all other common changes were present in the lineage that led to the majority of young insertions (Figure 6). Among the earliest inferred changes were premature stops in *gag* (CfERV-Fc1_CON_ bp 882 and 2203, respectively) and *env* (CfERV-Fc1_CON_ bp 6240) that tended to have been present among elements within a *Canis*-specific subclade. Additional inherited changes were present in a separate *Canis*-specific subclade in the form of a third distinct stop in *gag* (CfERV-Fc1_CON_ bp 1911). The shared *gag* stop was only observed within that cluster, suggesting its origin in a timeframe near variants contributing to the subclade followed by spread to new insertions therein. The stop is present in the chr17:9,744,973 and chr33:22,146,581 proviruses, therefore limiting LTR dating of the change; based on its restriction to assayed *Canis* members it likely originated within the last 2.74 my (Koepfli et al., 2015). Taken together, the data are consistent with independent origin and spread of multiple defective features that began prior to ancestors of the dog-like foxes and followed the Old and New World wolf split. The phylogenetic placement of defective proviruses suggests the co-occurrence of spread from multiple sources.

The apparent absence of any infectious retrovirus among canines is peculiar, particularly as individuals are likely to be challenged from viruses infecting prey species. While there have been reports of retroviral activities and particles displaying characteristic γ-like features in canine leukemias and lymphomas (Ghernati et al., 2000, Modiano et al., 2005, Modiano et al., 1995, Onions, 1980, Perk et al., 1992, Safran et al., 1992, Tomley et al., 1983), those findings have not been substantiated. It is well-known that canine cell lines are permissive for replication of retroviruses that infect other host species including human (Fadel and Poeschla, 2011), a property possibly reflecting the loss of the antiviral factor TRIM5α in canines (Sawyer et al., 2007). A recent report confirmed transcriptional activity from at least one γ-like CfERV group (non-Fc1(a)) in canine tissues and cell lines (Tarlinton et al., 2013). We have also preliminarily demonstrated expression of CfERV-Fc1(a) proviruses in canine tissues and tumor-derived cell lines (Jarosz and Halo, unpublished data).

Expression of ERV groups has been associated with both normal physiology and disease in several animal models, including humans, based on patterns of ERV-derived products observed within associated tissues (reviewed in (Jern and Coffin, 2008)). However, the consequences of this expression are not always clear. It is known from animal studies that ERVs with similarity to human ERVs, including those with extant forms with replicative activity, as well as proteins derived from related ERV members, are capable of driving aberrant cellular proliferation, tumorigenesis, and inciting immune responses (Jern and Coffin, 2008). Given our findings of the breadth and relative intactness of the CfERV-Fc1(a) lineage, we suggest that de-regulated expression from these loci is responsible for the γ-retroviral activities previously reported in canine tumors and cell lines, implying the potential for a pathogenic role of ERV-Fc1(a) loci and exogenous retroviruses in canines.

## Materials and Methods

### Whole genome sequence data

For ERV discovery, Illumina WGS data were obtained from a total of 101 samples corresponding to 37 breed dogs, 45 village dogs, and 19 wild canids (Auton et al., 2013, Botigue et al., 2017, Decker et al., 2015, Fan et al., 2016, Freedman et al., 2014, Marsden et al., 2016, Pendleton et al., 2018, Koepfli et al., 2015) (Table S1). Data were downloaded in fastq format and processed to Binary Alignment/Map BAM format using bwa version 7.15 and Picard v 2.9.0. SNV genotypes of sequenced samples were determined using Genome Analysis Toolkit (GATK) version 3.7 (McKenna et al., 2010). Information corresponding to all samples and sources of raw data is detailed in Table S1.

### Identification of annotated CfERVF1 reference insertions

The dog ERV-Fc1(a) lineage is classified in Repbase as ‘CfERVF1’ derived (Repbase update 10.08) (Jurka et al., 2005). We therefore mined the CanFam3.1 RepeatMasker output for elements classified as ‘CfERVF1_LTR’ and ‘CfERVF1-int’ according to Repbase vouchers to identify dog ERV-Fc1(a) LTRs and proviral elements, respectively. We required the presence of at least one LTR and contiguous internal sequence for a provirus, and the absence of any proximal internal region for a solo LTR. A total of 136 insertions were identified, corresponding to 21 proviral elements and 115 solo LTRs. The integration breakpoint ±1kb of each locus was extracted and used in BLAT searches against the other available carnivoran reference assemblies corresponding to ferret (MusPutFur1.0) (Peng et al., 2014), panda (BGI_Shenzhen1.0) (Li et al., 2010), and cat (Felis_catus_8.0) (Pontius et al., 2007) to confirm specificity to the dog reference. Sequences for proviral loci were extracted from CanFam3.1 based on the start and end positions of the full-length insertions, and filtered to remove severely truncated elements, resulting in 11 CfERV-Fc1(a) full-length or near full-length elements (*i.e.*, containing at least one viral gene region and associated 5’ or 3’ LTR). This count is consistent with recent findings of this ERV group in the dog reference (Diehl et al., 2016). Solo LTR insertions were filtered similarly to remove truncated elements, resulting in 96 insertions for further analysis.

### Deletion analysis of reference CfERV-Fc1(a) insertions

Reference insertions corresponding to deletion variants were inferred using the program Delly (v0.6.7) (Rausch et al., 2012), which processed BAM alignment files from samples indicated in Table S1 using a MAD score cutoff equal to 7, and a minimum map quality score threshold of at least 20. Resulting reference deletions with precise breakpoint predictions were next intersected with ‘CfERVF1’ reference coordinates based on RepeatMasker annotations of CanFam3.1. Only deletion calls corresponding to sizes of a solo LTR (400-500 bp) or a full-length provirus (7-9 kb) were considered for further analysis.

### Identification of non-reference of CfERV-Fc1(a) insertions

LTR-genome junctions corresponding to non-reference variants were assembled from supporting Illumina reads (Wildschutte et al., 2015, Wildschutte et al., 2016), with modifications as follows. The chromosomal positions of candidate non-reference ERVs were first identified using the program RetroSeq (Keane et al., 2013). Individual BAM files were queried using RetroSeq discovery to identify ERV-supporting discordant read pairs with one read aligned to the sequences corresponding to ‘CfERVF1’ and ‘CfERVF1_LTR’ from RepBase (Jurka et al., 2005). Individual BAM files were merged for subsequent steps using GATK as described (Wildschutte et al., 2016). RetroSeq call was run on the merged BAM files requiring ≥2 supporting read pairs for a call and output calls of levels 6, 7, and 8 further assessed, resulting in 2,381 candidate insertions. Output calls within ±500 bp of an annotated CfERV from the above queried classes were excluded to eliminate false calls of known loci. ERV-supporting read pairs and split reads within a 200 bp window of the call breakpoint were subjected to *de novo* assembly using the program CAP3 (Huang and Madan, 1999). Output contigs were filtered to identify ERV-genome junctions requiring ≥30 bp of assembled LTR-derived and genomic sequence in the form of (i) one LTR-genome junction, (ii) linked assemblies of 5’ and 3’ LTR junctions, or (ii) a fully resolved LTR (∼457bp) with clear breakpoints that mapped to CanFam3.1. Contigs that contained putative CfERV junctions were then aligned back to the reference to precisely map the insertion position of each call. Assembly comparisons were visualized using the program Miropeats (Parsons, 1995).

### Validations and allele screening

For validating non-reference calls, primers were designed to flank the predicted insertion within ∼200 bp based on the breakpoint position for a given site. Genomic DNA from a subset of samples with predicted insertion variants was used for validations. DNA with limited material was subjected to whole genome amplification (WGA) from ∼10ng genomic DNA according to the manufacturer’s protocol (Repli-G, Qiagen). For each sample, WGA DNA was diluted 1:20 in nuclease free water and 1 uL was utilized per PCR reaction. Two PCR reactions were run for each site in standard conditions using Taq polymerase (Invitrogen): one reaction utilized primers flanking each candidate call to detect the empty or solo LTR alleles; the second was to detect the presence of a proviral junction, utilizing the appropriate flanking primer paired with a primer within the CfERV-Fc1(a) proviral 5’UTR (near base ∼506 from the start of the Repbase F1 consensus element). Sanger sequencing was performed on at least one positive sample. When detected, provirus insertions were amplified in overlapping fragments from a single sample in a Picomaxx reaction per the manufacturer’s instructions (Stratagene) and sequenced to ≥4x across the full element. A consensus was then constructed for each insertion based on the Sanger reads obtained from each site. All sequences corresponding to non-reference solo-LTR insertions and all sequenced proviral elements have been made available in Table S2.

### Genomic distribution

The positions of the reference and non-reference insertions were intersected with Ensembl dog gene models (Release 81; ftp.ensembl.org/pub/release-81/gtf/canis_familiaris/). Intersections were performed using bedtools (Quinlan, 2014) with window sizes of 0, 5, 10, 25, 50, and 100 kb. To assess significant enrichment of insertions relative to genic regions, we performed one thousand permutations of randomly shuffled insertion positions, intersected the new positions with genes, and calculated the number of insertions intersecting genes within the varying window sizes as above. P-values were calculated as the number of permuted insertion sets out of one thousand that intersected with less than or equal to the number of genes observed in the true insertion set.

### Dating of individual proviruses

A molecular clock analysis based on LTR divergence was used to estimate times of insertion (Diehl et al., 2016, Johnson and Coffin, 1999, Wildschutte et al., 2016). For 7 non-reference and 8 reference proviruses that had 5’ and 3’ LTRs present, the nucleotide differences between those LTRs was calculated, treating gaps >2bp as single changes. The total number of changes was then divided by the LTR length (*e.g*. 457 bp), and the percent divergence normalized to the inferred canine background mutation rate of 1.3×10^−9^ changes per site per year (Botigue et al., 2017) to obtain age estimations in millions of years for individual insertions. The provirus at chr17:97,449,73 was excluded from the analysis due to truncation of its 3’ LTR. We extended LTR dating to estimate times of formation for identical LTR groups that included solo LTRs using a modification of the above approach as described elsewhere (Ishida et al., 2015). Briefly, the total length in bp of the LTRs making up each cluster was collectively added and the age estimate obtained by the percent divergence for a single base pair to have been introduced along the total length utilizing the same mutation rate of 1.3×10^−9^ changes per site per year.

### *In silico* genotyping

We genotyped 145 insertions (89 reference and 56 non-reference insertions) utilizing whole genome Illumina reads and reconstructed alleles corresponding to the empty and occupied sites. Genotyping was performed on 332 individuals including the 101 samples utilized for discoveries of polymorphic variants (Kim et al., 2012, Vamathevan et al., 2013, Owczarek-Lipska et al., 2013, Wang et al., 2013, Kim et al., 2013, Auton et al., 2013, Koepfli et al., 2015, Botigue et al., 2017, Freedman et al., 2014, Li et al., 2014, Zhang et al., 2014, Decker et al., 2015, Wang et al., 2016, Fan et al., 2016, Marsden et al., 2016, Robinson et al., 2016, Liu et al., 2014) (Table S4). Reference insertions were deemed to be suitable for genotyping based on manual assessment for the presence of paired TSDs and uninterrupted flanking sequence. Sites associated with duplication events were identified by comparison of flanking regions and TSD presence, and insertions within encompassing duplication (proviruses at chr3:219,396 and chrUn_JH373247:11,035), or situated within duplicated pre-insertion segments (chrUn_AAEX03025486:2,349) were excluded, as were sites with single assembled junctions (chr13:20,887,612; chr27:44,066,943; Table S2). The sequences from validated and completely assembled LTRs were utilized for allele reconstruction of non-reference sites. For example, the validated sequences for the non-reference solo LTRs at chr2:32,863,024 (8 bp LTR extension) and chr32:7,493,322 (associated with deletion of reference sequence) were included for genotyping of alternate alleles. For sites with linked, but non-resolved, 5’ and 3’ assembled junctions (*i.e*., missing internal sequence), we substituted the internal portion of each element from the Repbase CfERVF1 consensus (see Table S2), and used the inferred sequence for allele reconstruction. Insertion and pre-insertion alleles were then recreated based on ±600bp flanking each insertion point relative to the CanFam3.1 reference, accounting for each 5bp TSD pair. For each sample, genotype likelihoods were then assessed at each site based on re-mapping of those reads to either allele, with error probabilities based on read mapping quality (Li, 2011, Wildschutte et al., 2015), excluding sites without re-mapped reads for a given sample. Read pairs for which both reads mapped to the internal portion of the element were excluded to avoid false positive calls potentially introduced by non-specific alignment. The pipeline for genotyping is available at https://github.com/KiddLab/insertion-genotype. The genotyped samples were sorted by ancestral population, and allele frequencies estimated for the total number of individuals per population genotyped at each locus (Table S5).

### Admixture

A sample set containing only dogs and wolves were previously genotyped at approximately 7.6 million SNPs determined to capture genetic diversity across canids (Botigue et al., 2017). Using Plink (Purcell et al., 2007), sites were filtered to remove those with missing genotypes in at least ten percent of samples, those in LD with another SNP within 50 bp (--indep-pairwise 50 10 0.1), and randomly thinned to 500,000 SNPs. To reduce the bias of relatedness, the sample set was further filtered to remove duplicates within a single modern breed, leaving 254 samples (Table S7). Identification of wolf samples with high dog ancestry was made through five independent ADMIXTURE (Alexander et al., 2009) analyses of the thinned SNP set with random seeds (558905, 110684, 501738, 37781236, and 85140928) for K values 2 through 6. Since we aimed to discern cfERV-Fc1(a) insertions that may be dog-specific (*i.e.* having occurred since domestication), we removed any gray wolf that had high dog ancestry from further analysis. To do this, we calculated average dog ancestry within gray wolves at K=3 across all runs, which was the K value with the lowest cross validation error rate. Wolves with greater than 10% dog ancestry (an Israeli (isw01) and Spanish (spw01) wolf) were excluded from subsequent species and sub-population assessments.

### Phylogenetic analysis

Nucleotide alignments were performed using MUSCLE (Edgar, 2004) followed by manual editing in BioEdit (Hall, 1999) for intact CfERV-Fc1(a) LTRs from 19 proviral elements and 142 solo-LTRs. Of non-reference elements, the solo LTR with a 388 bp internal deletion at chr22:57,677,068 was excluded, as was the 141 bp truncated solo LTR at chr5:80,814,713. We also excluded partially reconstructed insertions corresponding to ‘one-sided’ assemblies or sites with linked 5’ and 3’ assembled junctions but that lacked internal resolution (Table S1). A maximum likelihood (ML) phylogeny was reconstructed from the LTR alignment using FastTree (Price et al., 2010) and the (GTR+CAT) model (generalized time reversible (GTR) model of nucleotide substitution plus “CAT” rate approximation). To infer the robustness of inferred splits in the phylogeny, local support values were calculated using the ML-based approach implemented in FastTree, wherein the Shimodaira-Hasegawa test is applied to the three alternate topologies (NNIs) around each node. The average d_N_/d_S_ ratio for intact env genes was determined using the codeml program in the PAML software package (version 4.8) (Xu and Yang, 2013) based on a Neighbor-Joining tree. Statistical significance was determined using the Nei-Gojobori method (Nei and Gojobori, 1986) implemented in MEGA7 (Kumar et al., 2016) with a null hypothesis of strict neutrality (d_N_ = d_S_).

## Acknowledgments

We thank John Coffin, Michael Freeman, Welkin Johnson and Zachary Williams for meaningful discussion and comments, and all owners and donors involved in sample donations for genomic DNA sources. We thank Anna Kukekova for sharing red fox genome data and Adam Boyko, Tomàs Marquès-Bonet, Carles Vilà, and Robert Wayne for early access to genome sequence data. Images of canids were obtained for *Urocyon littoralis* (“Island Fox II” (CC BY 2.0) by Shanthanu Bhardwaj), *Vulpes vulpes* (“El pequeño amigo” (CC BY 2.0) by Minette Lang), *Lycalopex culpaeus* (by Christian Mehlführer; Wikimedia Commons), *Cuon alpinus* (Wikimedia Commons), and *Canis lupus* (www.usda.gov). This work was supported in part by a National Institutes of Health Academic Research Enhancement Award R15GM122028 to JVH, National Institutes of Health grant R01GM103961 to JMK, National Institutes of Health Training Fellowship T32HG00040 to ALP, and UK Medical Research Council MC_UU_12014/10 to RJG. DNA samples were provided by the Cornell Veterinary Biobank, a resource built with the support of NIH grant R24GM082910 and the Cornell University College of Veterinary Medicine.

## Author Contributions

JVH, ALP, and JMK designed the study. JVH, ALP, and JMK were responsible for genome data processing. JVH, ASJ, MLD were responsible for sequence-based analysis. JVH, ALP, RJG and JMK were responsible for data analysis. JVH, ALP, and JMK wrote the paper. All authors have read and approved the final manuscript.

## Supplemental Figure Legends

**Figure S1. Assembled CfERV breakpoints remapped to the CanFam3.1 reference.** Three-way alignments for 58 non-reference insertions are shown. Alignments were used to depict CfERV-Fc1(a) LTR junctions obtained by assembled supporting reads (shown in red text) remapped to the CanFam3.1 reference sequence (shown in black text and underlined). The 5bp sequence corresponding to the target site duplication is underlined and bolded in the reference allele. The coordinates of the CanFam3.1 reference sequence shown is provided above each alignment; the first base of the LTR is labeled and indicated by an asterisk shown respective of orientation (‘+’ or ‘–’). Insertions for which a provirus was validated are labeled as appropriate. The single assembled junctions are provided for either of two insertions: chr13:20,998,612 (3’ junction); chr27:44,066,943 (5’ junction).

**Figure S2. Depletion of CfERV-Fc1(a) insertions near dog gene models.** Following one thousand permutations, the number of gene models that intersect with shuffled CfERV-Fc1(a) insertions are displayed in histograms. Permuted insertions that intersect with at least one Ensembl dog gene model precisely (green), within 10 kb (blue) or 50 kb (gray) are shown. Red lines indicate the observed number of insertions from the true set.

**Figure S3. Annotated CfERV-Fc1(a) consensus provirus.** A consensus provirus was deduced from 19 proviruses using BioEdit (http://www.mbio.ncsu.edu/BioEdit/bioedit.html) based on the most commonly represented nucleotide at each site. The consensus nucleotide sequence is shown in black text. The 5’ and 3’ LTRs are labeled with black bars. The translated sequences for the viral genes are indicated below and with bars at the right, with the Gag sequence in blue, Pol in orange, and Env in green. Motifs pertaining to viral functions are labeled appropriately on their translated sequence and general annotated in the right sidebar. Translated start and stop sites are indicated for each of the three genes. Segments for a predicted fusion peptide, membrane-anchoring TM region, and immunosuppressive domain (ISD) were determined using the program Phobius (http://phobius.sbc.su.se). Putative major splice donor and acceptor sites were determined using the program NetGene2 (http://www.cbs.dtu.dk/services/NetGene2/).

## Supplemental Tables

**Table S1. Canine sample information for discovery of CfERV-Fc1(a) insertions.** Information for the resequencing dataset of 101 canines used for CfERV-Fc1(a) insertion discovery. The sample identifier, sex, breed/species/population information and canine group is given per sample. Also provided are the Short Read Archive (SRA) sequence identifiers (SRR) matching the files downloaded and processed in this study, along with the PubMed identifier for the accompanying published study (if available) for each sample.

**Table S2. Information for non-reference sites considered in analyses.** The coordinates relative to CanFam3.1 are provided for each identified non-reference insertion. For each site, information pertaining to the insertion orientation, target site duplication (relative to the CanFam3.1 reference), detected insertion alleles (provirus, solo LTR), and element sequence is provided. Primer sequences are provided for validated sites. (A) Information for sequenced loci and validated sequences. (B) Information for loci with complete assembled insertion alleles. (C) Information for loci with partially assembled insertion alleles.

**Table S3. Gene region information and GO ontology analyses.** The coordinates for each reference and non-reference insertion are provided along with Ensembl gene models from dog (release #81) that are within window distances of 0, 5, 10, 25, 50, and 100 kb of the insertion.

**Table S4. Sample information for canid genotyping.** Sample and data access information for the resequencing dataset of 332 canines genotyped at the discovered CfERV-Fc1(a) reference and non-reference insertions. Accompanying data descriptions provided for each sample match that of Table S1.

**Table S5. Genotypes and inferred allele frequencies.** Raw genotypes obtained across 332 resequenced samples for 56 non-reference and 89 reference insertions are provided in vcf format. Allele frequencies were calculated from raw genotypes per canid species or sub-population, as indicated above each column. Non-genotyped sites are noted with an “-”.

**Table S6. LTR nucleotide alignment.** LTR alignment for phylogenetic analysis using LTRs from a total of 19 proviruses and 142 solo LTRs, provided in fasta format.

**Table S7. Samples included in admixture analysis.** Sample information for the 254 samples included in admixture analysis. Accompanying data columns provided for each sample match that of Table S1.

